# A bacterial CARD-NLR immune system controls the release of gene transfer agents

**DOI:** 10.1101/2025.05.08.652646

**Authors:** Emma J. Banks, Pavol Bárdy, Ngat T. Tran, Phuong M. Nguyen, Abbas Maqbool, Tung B. K. Le

## Abstract

Bacteria have evolved a wide array of immune systems to detect and defend against external threats including mobile genetic elements (MGEs) such as bacteriophages, plasmids, and transposons. MGEs are often selfish, exploiting their bacterial hosts to propagate, however they can also provide adaptive advantages through horizontal gene transfer. Gene transfer agents (GTAs), which are non-infectious domesticated prophages, represent a unique class of beneficial MGEs that facilitate bacterial gene transfer. Despite their domestication, GTAs retain phage-like features, including the requirement for host cell lysis to release particles, that may inadvertently trigger host immunity. How GTAs might avoid, subvert, or possibly adopt host immune systems to complete their life stages is poorly understood. Here, we identify a tripartite system, LypABC, that is essential for GTA-mediated cell lysis in *Caulobacter crescentus*. LypABC resembles caspase recruitment domain-nucleotide-binding leucine-rich repeat (CARD-NLR) anti-phage defence systems that mediate abortive infection wherein infected cells die to prevent phage proliferation, thereby protecting the overall bacterial population. LypABC-deficient cells produce host DNA-packed GTA particles and eventually die but cannot lyse to release GTA particles. Moreover, overproduction of LypABC is highly toxic to both GTA-producing and non-producing cells, highlighting the need for strict regulation. We find that such regulation is achieved transcriptionally by a repressor, RogB, which binds the promoters of *lypABC* and of essential GTA activator genes, thus coupling GTA activation and host cell lysis. While traditionally considered antagonistic towards MGEs, our findings here suggest that immunity components are versatile and can be adapted to support MGEs.

## INTRODUCTION

Mobile genetic elements (MGEs), such as bacteriophages, plasmids, and transposons, are double-edged swords; while they can confer certain adaptive advantages to their host through horizontal gene transfer, they often act selfishly, exploiting the host for their own propagation (*1*). Bacteria are therefore engaged in an arms race against MGEs and have evolved an extraordinary diversity of immune systems to detect and defend against MGEs, including 150 distinct anti-MGE systems that have been identified in recent years (*2–9*). Although immune systems are traditionally considered antagonistic to MGEs, it remains unclear whether some immune systems might be versatile and, in certain contexts, may facilitate rather than prevent MGE propagation.

Amid the constant conflict between bacteria and phages, gene transfer agents (GTAs) are exceptions. GTAs are selfless virus-like MGEs which have been domesticated from ancient viruses to provide beneficial functions to their hosts (*10–12*). GTAs (*13–15*) are encoded by gene clusters within a wide variety of bacterial and archaeal genomes, and are deeply integrated with their host physiology (*16, 17*). GTAs transition through a series of life stages: GTA gene cluster activation (*18*), GTA particle assembly (*19*), non-selective encapsulation of host DNA into GTA particles (*20, 21*), GTA particle release by host cell lysis (*22, 23*), and finally transfer of host DNA into recipient bacteria (*19, 24, 25*). Most notably, unlike bacteriophages, GTA capsid heads are too small to package complete GTA gene clusters (for example, the *Caulobacter crescentus* GTA can package only ∼8.3 kb of DNA yet its encoding GTA cluster is >15 kb (*26*)). Consequently, GTAs are unable to self-multiply and be infectious (*27*). Despite GTA domestication, the phage-like origin and appearance of GTAs - together with life stages that include host cell lysis - are factors that may inadvertently trigger host immunity. It remains unclear how GTAs might avoid, subvert, or even adopt host immune systems to complete their life stages.

Here, by studying GTA-mediated host cell lysis in *C. crescentus*, we identified a potential bacterial immune system that has been adopted to control the release of GTA particles. The immunity genes resemble components of caspase recruitment domain-nucleotide-binding leucine-rich repeat (CARD-NLR) anti-phage defence systems (*28*). First described in *Lysobacter enzymogenes*, CARD-NLR immunity occurs through abortive infection which involves sacrificial death of phage-infected cells, preventing the release of mature phage particles, and thereby curbing infection (*28*). The *L. enzymogenes* CARD-NLR system senses phage infection, somehow activating a CARD-containing protein component which interacts with an NLR-like protein (*28*). Cell death occurs through proteolysis-based activation of a gasdermin effector which directly causes cell lysis by forming membrane pores and permeabilising the cell membrane (*29, 30*). In animals, many NLR-based inflammatory responses also contain CARD components that signal to caspases (*31, 32*) which then proteolytically cleave and activate gasdermin effectors, leading to the release of pro-inflammatory cytokines and cell death (*33–35*).

We find that GTA-mediated lysis in *C. crescentus* occurs via the action of LypABC which resemble a CARD-NLR anti-phage immune system. Mutagenesis of *lypABC* revealed that predicted anti-phage defence domains are essential for cell lysis. We further show that in the absence of *lypABC*, host cell lysis is abolished without affecting GTA particle production or host cell death, indicating that LypABC specifically mediate cell lysis for GTA release. AlphaFold-Multimer and bacterial two-hybrid assays indicate that LypA and LypB form a co-complex mediated by the CARD-like domain of LypA. Overproduction of LypABC is highly toxic to both GTA-producing and non-producing cells, highlighting the need for this system to be tightly regulated. Lastly, we identify a transcriptional regulator, RogB, that directly represses genes encoding GTA-activating factors and LypABC, thereby coupling GTA gene cluster activation and host cell lysis. In summary, we present evidence that immune systems are versatile and can perform new biological functions that are beneficial, rather than antagonistic, to MGEs.

## RESULTS

### GTA-mediated host cell lysis results in ghost cell formation

*C. crescentus* GTA synthesis is repressed under standard laboratory conditions but can be activated by deleting the master repressor gene, *rogA* (*26*) (**Figure 1A**). This relieves RogA-mediated repression of the *gafYZ* operon which is essential for GTA activation (*26*). The transcriptional activator GafY, together with integration host factor (IHF), co-activates the expression of GTA gene clusters and accessory genes elsewhere on the chromosome. Meanwhile, GafZ enables RNA polymerase to bypass internal transcription terminators within the core GTA gene cluster, ensuring complete expression of an entire biosynthetic gene cluster (*26, 36, 37*) (**Figure 1A**).

**Figure 1.**
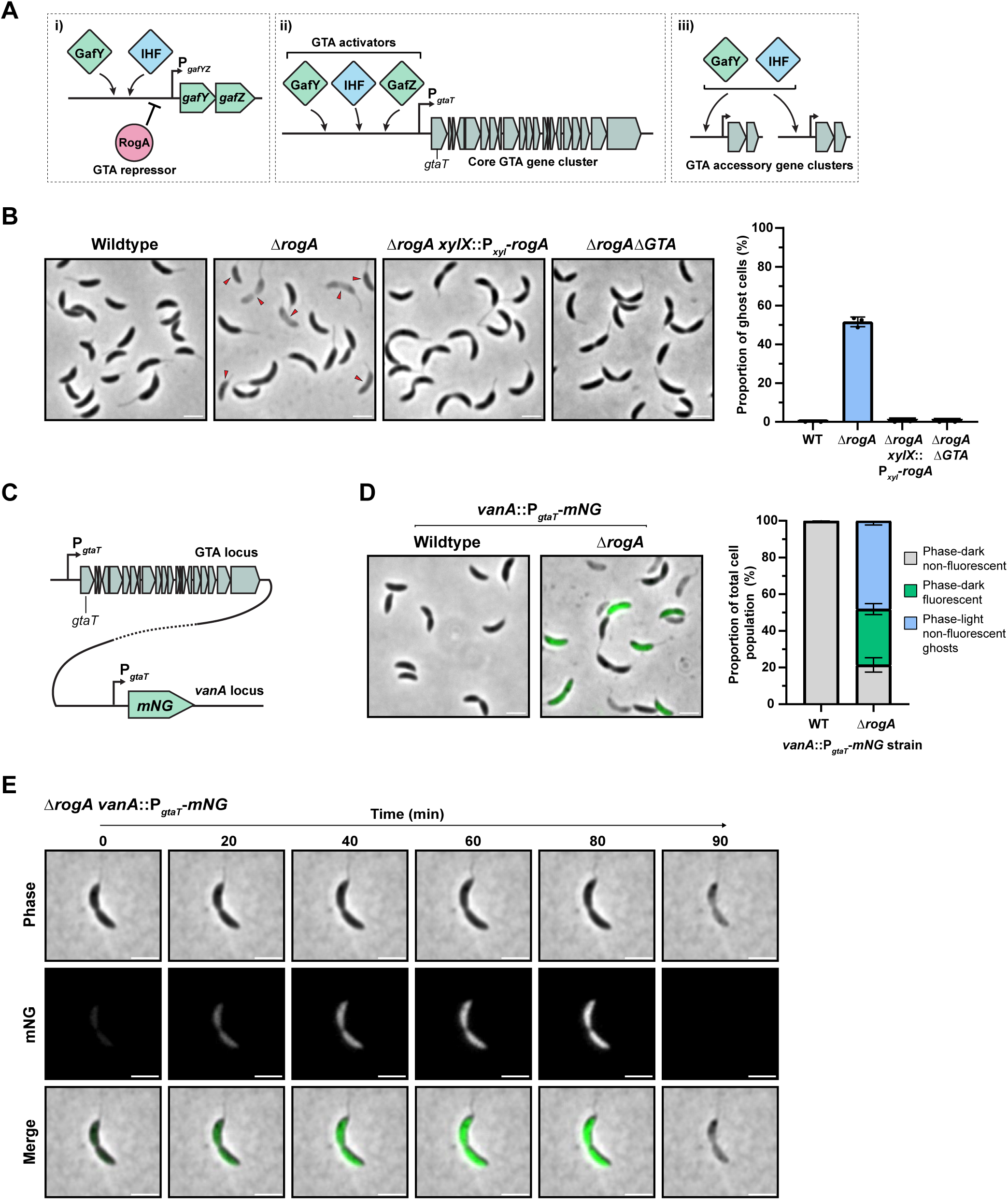
GTA-mediated host cell lysis results in ghost cell formation. **(A)** Schematic of gene transfer agent (GTA) regulation in *Caulobacter crescentus*. i) GafY and IHF positively activate transcription from the *gafYZ* locus, however the GTA repressor protein RogA can repress transcription of *gafYZ*. ii) GTA activating proteins GafY, IHF and GafZ bind to the core GTA gene cluster promoter, P*_gtaT_*, and activate cluster transcription, resulting in GTA activation. iii) GafY and IHF also activate transcription from the promoters of accessory GTA gene clusters at different chromosomal loci. **(B)** Left: *C. crescentus* strains visualised by phase-contrast microscopy. The Δ*rogA* mutant was complemented by expression of *rogA* from the *C. crescentus* chromosomal *xylX* locus promoter (P*_xyl_*). Gene expression was induced by the addition of 0.3% xylose to growth media. Images are representative of three independent repeats. Red arrows indicate ghost cells. Scale bar: 2 µm. Right: quantification of the proportion of ghost cells within the population of each strain. Data represent the mean ±. s.d from three independent experiments. n=400 cells analysed per experiment. **(C)** Schematic of strain construction for an mNeonGreen (mNG) reporter for GTA activation. The core GTA gene cluster was unaltered and a copy of the P*_gtaT_* promoter inserted at the neutral *vanA* locus, immediately followed by the *mNG* gene. The dashed line indicates the large genomic distance separating the two loci **(D)** Left: fluorescence microscopy images of wildtype and Δ*rogA* strains containing the *vanA*::P*_gtaT_* -*mNG* reporter fusion. Images are representative of three independent repeats. Scale bar: 2 µm. Right: quantification of three different cell types within each population: phase-dark non-fluorescent cells (grey bar), phase-dark fluorescent cells (green bar), and phase-light non-fluorescent ghost cells (blue bar). Data represent the mean ±. s.d from three independent experiments. n=400 cells analysed per experiment. **(E)** Fluorescence microscopy images from a time-lapse experiment performed with the Δ*rogA vanA::*P*_gtaT_* -*mNG* strain showing GTA activation (mNG signal) followed by cell lysis. The time-lapse can be viewed in Movie S1. Additional examples are presented in Figure S1, and Movies S2-3. Data are representative of at least three independent repeats. Scale bar: 2 µm.

To investigate the consequences of GTA activation and how this leads to host cell lysis, we observed wildtype (GTA-off) and Δ*rogA* (GTA-on) *C. crescentus* strains during stationary phase by phase-contrast microscopy (**Figure 1B**). While the wildtype strain comprised almost entirely phase-dark cells with only 0.1 ± 0.1% phase-light ghost cells, the Δ*rogA* mutant population was heterogeneous, consisting of a mixture of phase-dark cells and a significantly higher proportion of phase-light ghost cells (51.6 ± 2.5%) than the wildtype. Complementation of the Δ*rogA* mutant restored the wildtype phenotype. Furthermore, deletion of the entire core GTA gene cluster in the Δ*rogA* background (Δ*rogA*Δ*GTA*) completely eliminated ghost cell formation, indicating that GTA activation resulted in the observed ghost cells i.e. remnants of *C. crescentus* cells that have undergone lysis (**Figure 1B**).

Next, to monitor GTA activity from cluster activation to host cell lysis at the single-cell level, we engineered a fluorescent reporter by transcriptionally fusing the promoter of the core GTA cluster (P*_gtaT_*) to mNeonGreen (mNG) and integrated this construct ectopically at the vanillate-utilisation (*vanA*) locus (**Figure 1C**). As expected, no mNG-fluorescent cells were observed in the wildtype strain. In contrast, the Δ*rogA* mutant showed a mixture of mNG-fluorescent cells (30.4 ± 3.0%) and non-fluorescent cells (of which 21.5 ± 3.9% were phase-dark and 48.2 ± 2.3% were phase-light ghost cells) (**Figure 1D**). Subsequent time-lapse microscopy revealed the emergence of mNG signal, followed by cell death within ∼90 minutes. Cell death typically involved the loss of cytoplasmic mNG signal, cell pole contraction, and then a transition from a phase-dark to phase-light ghost cell state - all within ∼10 minutes (**Figure 1E, Figure S1, Movies S1-3**). Notably, ghost cells maintained their vibrioid morphology, contrasting with modes of phage holin/endolysin-mediated lysis that often involve outer membrane blebbing and morphological deformation, culminating in explosive cell lysis (*38–42*). Altogether, these observations highlight the heterogeneous nature of GTA production in *C. crescentus* and suggest that GTAs may cause host cell lysis via a mechanism distinct from classical holin-endolysin pathways.

### The LypABC system is essential for GTA-mediated host lysis

Consistent with the observed non-explosive GTA-mediated lysis phenotype, by bioinformatic searches, we found no homologues of canonical holin/endolysin-encoding genes within the core GTA cluster or elsewhere on the *C. crescentus* genome. To identify candidate lysis genes in an unbiased manner, we conducted saturated transposon mutagenesis combined with deep sequencing (Tn-seq), comparing transposon insertion frequencies between wildtype (GTA-off) and Δ*rogA* (GTA-on) strains (**Figure 2A** and **Table S1**). We reasoned that transposon insertions disrupting either GTA activator-encoding genes or genes required for host cell lysis would prevent lysis, leading to a higher recovery of DNA for deep sequencing. Accordingly, we anticipated a higher frequency of insertions within genes required for GTA activation or cell lysis in Δ*rogA* compared to the wildtype background.

**Figure 2.**
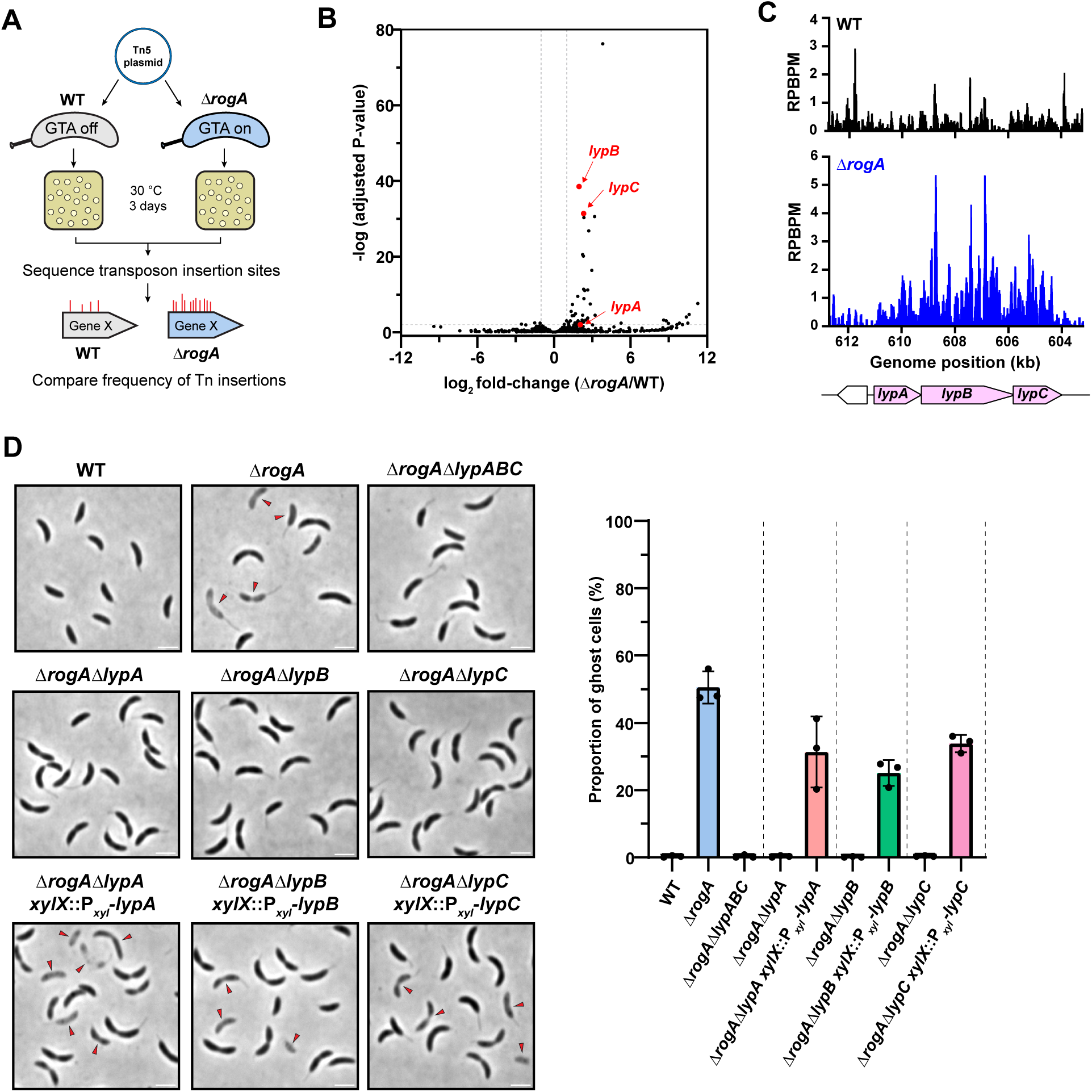
The LypABC system is essential for GTA-mediated host lysis. **(A)** Schematic of Tn-seq transposon mutagenesis method to identify potential lysis genes. **(B)** Tn-seq volcano plot showing the log_2_ fold-change of transposon insertions comparing the Δ*rogA* strain to the wildtype strain. Grey lines indicate a value of 2.0. *lypA*, *lypB*, and *lypC*, which contain more transposons in the Δ*rogA* strain are annotated.Three independent repeats were performed. **(C)** Plots showing the reads per base pair per million reads (RPBPM) at the *lypABC* genomic locus, indicating the higher frequency of transposon insertions throughout the *lypABC* genes within the Δ*rogA* strain (bottom, blue trace) compared to the wildtype strain (top, black trace). **(D)** Left: phase-contrast microscopy images of each Δ*rogA* Δ*lyp* gene deletion mutant. Deletion mutants were complemented by expressing the appropriate *lyp* gene from the *C. crescentus* chromosomal *xylX* locus promoter (P*_xyl_*). Gene expression was induced by the addition of 0.3% xylose to growth media. Images are representative of three independent repeats. Red arrows indicate ghost cells. Scale bar: 2 µm. Right: quantification of the proportion of ghost cells within the population of each strain. Data represent the mean ±. s.d from three independent experiments. n=400 cells analysed per experiment.

Our Tn-seq libraries achieved near-saturating coverage, with an average of one insertion every ∼10-20 bp. As expected, we found genes encoding known GTA activators, such as GafY, GafZ, and the α and β subunits of IHF, which showed 7.8-fold, 6.6-fold, 8.3-fold, and 4.9-fold increases in transposon insertions, respectively, confirming that the Tn-seq experiments worked (**Figure S2**). Differential analysis of the data identified 41 genes with significantly enriched Tn insertions in the Δ*rogA* background (log_2_ fold-change ≥ 1.0, adjusted p-value <0.01) (**Figure 2B**). Among these, we identified an operon comprising the three genes CCNA_03886, CCNA_00580, and CCNA_00579 (renamed to *lypABC*, respectively, for putative <u>ly</u>sis <u>p</u>roteins <u>ABC)</u> that contained higher insertion frequencies in the Δ*rogA* background compared to wildtype (4.2-fold, 3.9-fold, and 5.0-fold, respectively) (**Figure 2B** and **Figure 2C**). This 6.9 kb operon, located ∼2.4 Mb away from the GTA gene cluster, is predicted by the bioinformatic tool DefenseFinder (*43*) to encode an immune system resembling a CARD-NLR anti-phage defence complex. Deletion of either the entire *lypABC* operon or individual *lyp* genes in the Δ*rogA* background (GTA-on) completely abolished ghost cell formation (**Figure 2D**). Meanwhile, complementation of each Δ*rogA*Δ*lyp* mutant by expressing wildtype gene copies from an ectopic xylose-utilisation (*xylX*) locus restored the lytic phenotype (**Figure 2D**). These data demonstrate that all three *lyp* genes are required for GTA-mediated host cell lysis.

### LypABC are dispensable for DNA packaging and cell death but are essential for GTA particle release

To investigate whether LypABC are involved in the production of DNA-packaged GTA particles within *C. crescentus* cells, we assayed for the presence of a ∼8.3 kb DNA band in total DNA extractions (*26*). The DNA band was visible in the Δ*rogA*Δ*lypABC* mutant as well as in each of the individual Δ*rogA*Δ*lyp* gene mutants, indicating that LypABC are not required for the encapsulation of host genomic DNA into GTA particles (**Figure 3A**). To determine whether GTA particles are produced in the absence of *lypABC*, we performed immunoblotting using an antibody against the GTA head-tail connector structural protein, GtaL (*36*). While GtaL was detected in both the intracellular and extracellular fractions of the Δ*rogA* mutant, consistent with particle release via cell lysis, GtaL was present only in the intracellular fraction of the Δ*rogA*Δ*lypABC* mutant, suggesting that GTA particles assemble but are not released from cells to the extracellular space (**Figure 3B**). We further visualised the formation and release of GTA particles directly using cryo-electron tomography. GTA capsids were present in both Δ*rogA* and Δ*rogA*Δ*lypABC* cells, however, membrane collapse and subsequent cell lysis to release GTA particles only occurred in Δ*rogA* cells (**Figure 3C** and **Figure S3**), suggesting that LypABC are required for host lysis but not for upstream intracellular particle assembly.

**Figure 3.**
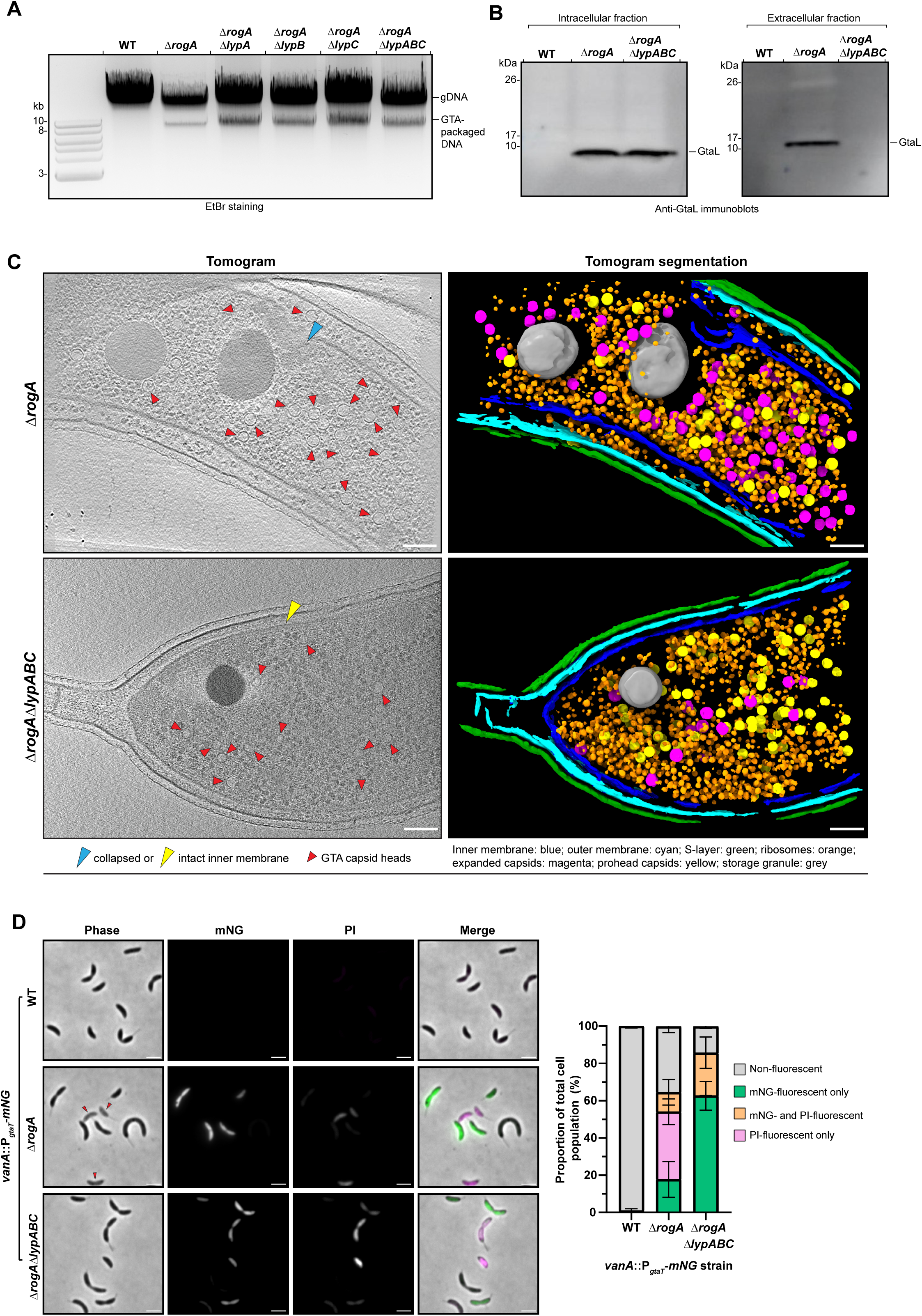
LypABC are dispensable for DNA packaging and cell death but are essential for GTA particle release. **(A)** Total genomic DNA extractions from *C. crescentus* strains visualised by agarose gel electrophoresis and ethidium bromide (EtBr) staining. GTA-packaging of bacterial host DNA is determined by presence of an 8.3 kb band. A representative gel from two independent experiments is presented. **(B)** Immunoblots of intracellular (left) and extracellular supernatant (right) fractions from *C. crescentus* strains using a polyclonal antibody targeting the GTA structural head-tail connector protein, GtaL. The immunoblot is representative of two independent experiments. **(C)** Cryo-electron tomograms of Δ*rogA* and Δ*rogA*Δ*lypABC* cells (left) with segmented tomograms for each image (right). Red arrows: GTA capsid heads, blue arrow: collapsed inner membrane, yellow arrow: intact inner membrane. Segmentation labels: inner membrane: blue; outer membrane: cyan; S-layer: green; ribosomes: orange; expanded capsids: magenta; prohead capsids: yellow; phosphate granule grey. Scale bar: 100 nm. Images are representative of two independent experiments. Additional images are shown in Figure S3. **(D)** Left: fluorescence microscopy images of *C. crescentus* strains containing a *vanA*::P*_gtaT_*-*mNG* fluorescent reporter for GTA activation. Cells were incubated with the dead cell-permeable dye propidium iodide (PI) and visualised on agarose pads. Individual fluorescent channels and the channel merge (green: mNG, magenta: PI) are shown. Images are representative of three independent repeats. Red arrows indicate ghost cells. Scale bar: 2 µm. Time-lapses of the Δ*rogA*Δ*lypABC vanA*::P*_gtaT_*-*mNG* strain grown in the presence of PI can be viewed in Figure S4 and Movies S4-5. Right: quantification of four different cell types within each population: non-fluorescent cells (grey bar), mNG-fluorescent only cells (green bar), mNG- and PI-fluorescent cells (orange bar), and PI-fluorescent only cells (pink bar). Data represent the mean ±. s.d from three independent experiments. n=400 cells analysed per experiment.

To investigate whether GTA-producing cells lacking LypABC are viable, we employed dual-labelling with propidium iodide (PI), a fluorescent DNA stain that can only enter membrane-compromised dead cells, and the P*_gtaT_*-mNG reporter. Wildtype cells were phase-dark with no detectable mNG or PI signal (**Figure 3D**), and - as observed previously (**Figure 1D)** - the Δ*rogA* population was heterogeneous, containing both mNG-fluorescent and non-fluorescent phase-dark cells in addition to non-fluorescent ghost cells. Weak PI signal was visible in most but not all Δ*rogA* ghost cells (36.3 ± 6.9%); PI fluoresces upon DNA binding, thus this weak PI signal may be due to the release of DNA-packed GTA particles. In the non-lysing Δ*rogA*Δ*lypABC* mutant, 85.7 ± 0.9% of all cells contained mNG signal, confirming that GTA cluster expression occurs independently of LypABC (**Figure 3D**). Importantly, 23.0 ± 8.5% of all cells exhibited both mNG and PI signal (**Figure 3D**), indicating that these GTA-producing cells were non-viable despite the absence of visible lysis. This conclusion was further corroborated by observing cells growing on PI-supplemented agarose pads by time-lapse microscopy (**Figure S4, Movies S4-5**). Similar to Δ*rogA* cells (**Figure 1E**), in Δ*rogA*Δ*lypABC* cells, mNG signal appeared first, followed cell pole contraction (**Figure S4, Movies S4-5**). However, unlike the Δ*rogA* strain, cells did not transition to a phase-light ghost cell state. Instead, the mNG signal was gradually replaced by PI signal over ∼500 minutes, suggesting that cells die in the absence of visible lysis (likely due to host genome digestion and packaging into GTA capsids). Thus far, the collective evidence suggests that LypABC are essential for host cell lysis to release GTA particles, but are dispensable for GTA gene expression, DNA packaging and GTA-mediated cell death.

### LypABC resemble a CARD-NLR anti-phage defence system

Bioinformatic analysis by DefenseFinder (*43*) suggested that LypABC may architecturally resemble a CARD-NLR anti-phage immune system (**Figure 4A**). First identified in *L. enzymogenes*, CARD-NLR systems mediate abortive infection by sensing phage invasion and then activating a death-domain effector (e.g. a pore-forming gasdermin protein in *L. enzymogenes*), leading to cell death and infection containment (*44*).

**Figure 4.**
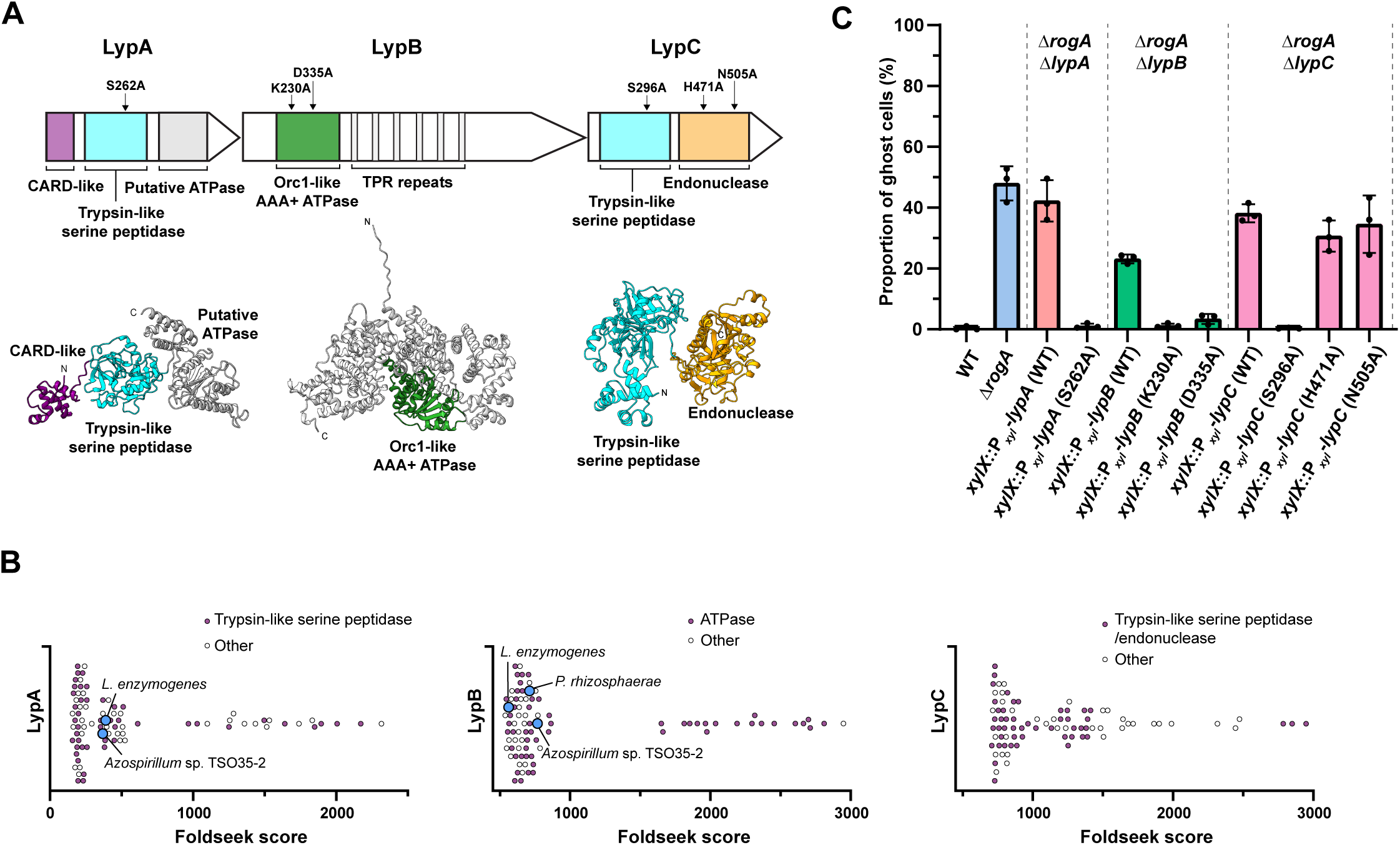
LypABC resemble a CARD-NLR anti-phage defence system. **(A)** Schematics showing the predicted domain organisation for each LypABC protein (top) with AlphaFold3-predicted structures (bottom). **(B)** Alpha-Fold-predicted structural homologues of LypA (left), LypB (middle), and LypC (right) identified by Foldseek with a query coverage >80% and E-value <5.88 x10^-9^, sorted according to Foldseek score. Blue circles: characterised CARD-NLR homologues; purple circles: homologues that share the same predicted catalytic domain as the query protein; white circles: homologues with a different domain annotation or annotated as uncharacterised (”other”). **(C)** Quantification of the proportion of ghost cells within the population of different Δ*rogA*Δ*lyp* mutants complemented with either the wildtype *lyp* gene or a *lyp* gene variant containing a point mutation of a predicted catalytic amino acid. The location of each amino acid mutation is indicated by arrows above the LypABC protein schematics in (A). Means ± standard deviations of three independent repeats are shown. n=400 cells analysed per repeat.

To explore the significance of LypABC’s resemblance to CARD-NLR systems, we performed further analysis and structural modelling for each LypABC component. Foldseek-based (*45*) searches for structural homologues of LypA and LypB returned known CARD-NLR proteins including the trypsin-like serine peptidase and NLR-like protein components of the *L. enzymogenes* and *Azospirillum* sp. CARD-NLR systems (*44*), while LypC returned trypsin-like serine peptidase and endonuclease homologues (**Figure 4B**), Moreover, AlphaFold3-generated models of LypA and LypB superimpose closely to their *L. enzymogenes* homologues (**Figure S5**), indicating that LypABC indeed belong to the CARD-NLR protein family.

LypA, a 592 amino acid (aa) protein, contains three predicted domains: an N-terminal α-helical bundle domain (aa 1-80), a middle trypsin-like serine peptidase domain (aa 85-305), and a C-terminal ATPase domain (aa 324-592). While structural modelling of LypA with ATP and Mg^2+^ by AlphaFold3 suggested potential ATP-binding ability (**Figure S6A**), canonical Walker A/B motifs are absent, thus we amended the annotation of this C-terminal domain to “putative ATPase”. The LypA N-terminal domain resembles eukaryotic caspase recruitment domains (CARDs) (*44*). However, while predicted to share structural similarity with the X-ray crystallography CARD structures of *L. enzymogenes* and *Azospirillum* sp., the AlphaFold3 model of the LypA N-terminal domain comprises only five α-helices in contrast to the six α-helices present in a canonical CARD (*46*) (**Figure S7**). In *L. enzymogenes*, the CARD is suggested to activate a gasdermin effector protein which causes cell lysis (*44*). To assess the relevance of the CARD-like domain in LypA, we generated a CARD-truncated *lypA* mutant, however the resulting truncated protein was unstable and undetectable *in vivo* (**Figure S8A**), preventing us from drawing further conclusions. Next, we assessed the role of the trypsin-like serine peptidase domain of LypA, discovering that expressing a *lypA* catalytic serine residue mutant (S262A) did not restore lysis in the Δ*rogA*Δ*lypA* background (**Figure 4C**) despite robust protein expression (**Figure S8A**). This indicates that the peptidase activity of LypA is critical for LypABC-mediated lysis.

LypB contains a predicted Orc1-like AAA+ ATPase domain (aa 193-392) containing conserved Walker A (Gx_4_GKS) and Walker B (MVLD) motifs, followed by an extended region of tandem tetratricopeptide (TPR)-like repeats (aa 393-771) (**Figure 4A, Figure S5B** and **Figure S9**). This domain architecture is reminiscent of bacterial NLR proteins. In addition to Foldseek (**Figure 4B**), HHPred searches further identified known bacterial NLR proteins as homologues of LypB including MalT from *E. coli* (*47*) and Avs3 from *Salmonella enterica* (*48*). NLR ATPase domains typically bind ATP and oligomerise into a large macromolecular structure (*49*). Expressing *lypB* variants with point mutations in the Walker A (i.e. K230A) and Walker B (i.e. D335A) motifs failed to complement the Δ*rogA*Δ*lypB* mutant phenotype (**Figure 4C**) despite wild-type levels of protein expression (**Figure 8B**), suggesting that the ATPase activity of LypB is essential for LypABC-mediated host cell lysis.

Considering that the trypsin-like serine peptidase protein and NLR-like protein of the *L. enzymogenes* anti-phage system interact (*44*), we tested whether LypA and LypB form a co-complex. AlphaFold3-Multimer predicted an interaction between the CARD of LypA and the N-terminal domain of LypB (ipTM score: 0.85) (**Figure S10A**). We validated this predicted interaction by performing *in vivo* bacterial two-hybrid experiments, demonstrating that the LypA CARD interacts solely with the N-terminal domain of LypB, but no other domain encoded by the LypABC system (**Figure S10C**). Moreover, the LypA CARD could self-interact (**Figure S10B** and **Figure S10C**), consistent with findings that eukaryotic CARDS often form homomeric interactions (*50–52*).

LypC is predicted to contain two distinct domains: an N-terminal trypsin-like serine peptidase domain (aa 1-335) and a C-terminal endonuclease domain (aa 354-662), separated by a flexible linker (**Figure 4A**). Using a similar complementation approach, we discovered that the serine peptidase activity of LypC was essential for host lysis, as assessed via the catalytic residue S296A mutation (**Figure 4C**). However, disrupting the endonuclease activity, via mutating catalytic residues H471 or N505 to alanine, had no effect on LypABC-mediated cell lysis (**Figure 4C**). Immunoblotting confirmed stable production of all mutant proteins (**Figure S8B**). The LypABC system does not encode a gasdermin, however in other CARD-NLR variants, gasdermins are replaced by alternative death-domain effectors such as phospholipases or endonucleases (*28*). While LypC encodes an endonuclease domain, our mutagenesis analysis suggests that this domain is not the death-domain effector in *C. crescentus*. Notably, the LypC homologue in *Caulobacter endophyticus* lacks an endonuclease domain entirely (**Figure S11A**), further suggesting that GTA-mediated cell lysis is endonuclease independent.

Unlike most anti-phage defence systems whose encoding genes often cluster into genomic “defence islands”, the *C. crescentus lypABC* operon is not associated with any known or predicted immunity genes (**Figure S11B**). Furthermore, CARD-NLR systems are rare in bacteria, present in only ∼0.35% of sequenced bacterial genomes, as surveyed by the DefenseFinder webservice (*43*). Indeed, *lypABC* is not conserved even in *Caulobacter* species; we could only detect complete *lypABC* operons in four of the eleven *Caulobacter* species predicted to also encode GTAs: *C. crescentus, C. radicis*, *C. zeae*, and *C. endophyticus* (**Figure S11C**). Together, these results suggest that *C. crescentus* may have evolved a new lysis pathway, based on an anti-phage defence system, to facilitate the release of GTA particles.

### Misregulation of *lypABC* expression is highly toxic to *C. crescentus*

Classical abortive infection systems must be tightly regulated to avoid self-inflicted cell death through untimely activation i.e. autoimmunity (*53*). To investigate whether misregulation of *lypABC* expression could result in such toxicity, we inserted a strong cumate-inducible promoter (*54*) immediately upstream of the *lypABC* operon, creating P*_cumate_*-*lypABC* strains in either the wildtype or the Δ*rogA* background (**Figure 5A**). Addition of cumate had no effect on the viability of wildtype cells nor did it cause an elevated proportion of ghost cells in the Δ*rogA* mutant (**Figure 5B**). In the Δ*rogA* P*_cumate_*-*lypABC* strain, there were 25.1 ± 8.9% ghost cells in the absence of cumate-induction - likely due to transcriptional bleed-through from the weaker upstream native P*_lypABC_* promoter. Strikingly, cumate-induction in this same strain led to a dramatic increase in ghost cells (89.0 ± 1.8% in total). A similarly high level of cell death (88.8 ± 5.5% ghost cells) was observed upon cumate-induction of P*_cumate_*-*lypABC* in the wildtype background (**Figure 5B**). These findings demonstrate that overexpression, and thus misregulation, of *lypABC* is highly toxic to both GTA-producing and non-producing cells, highlighting the need for the system to be tightly regulated.

**Figure 5.**
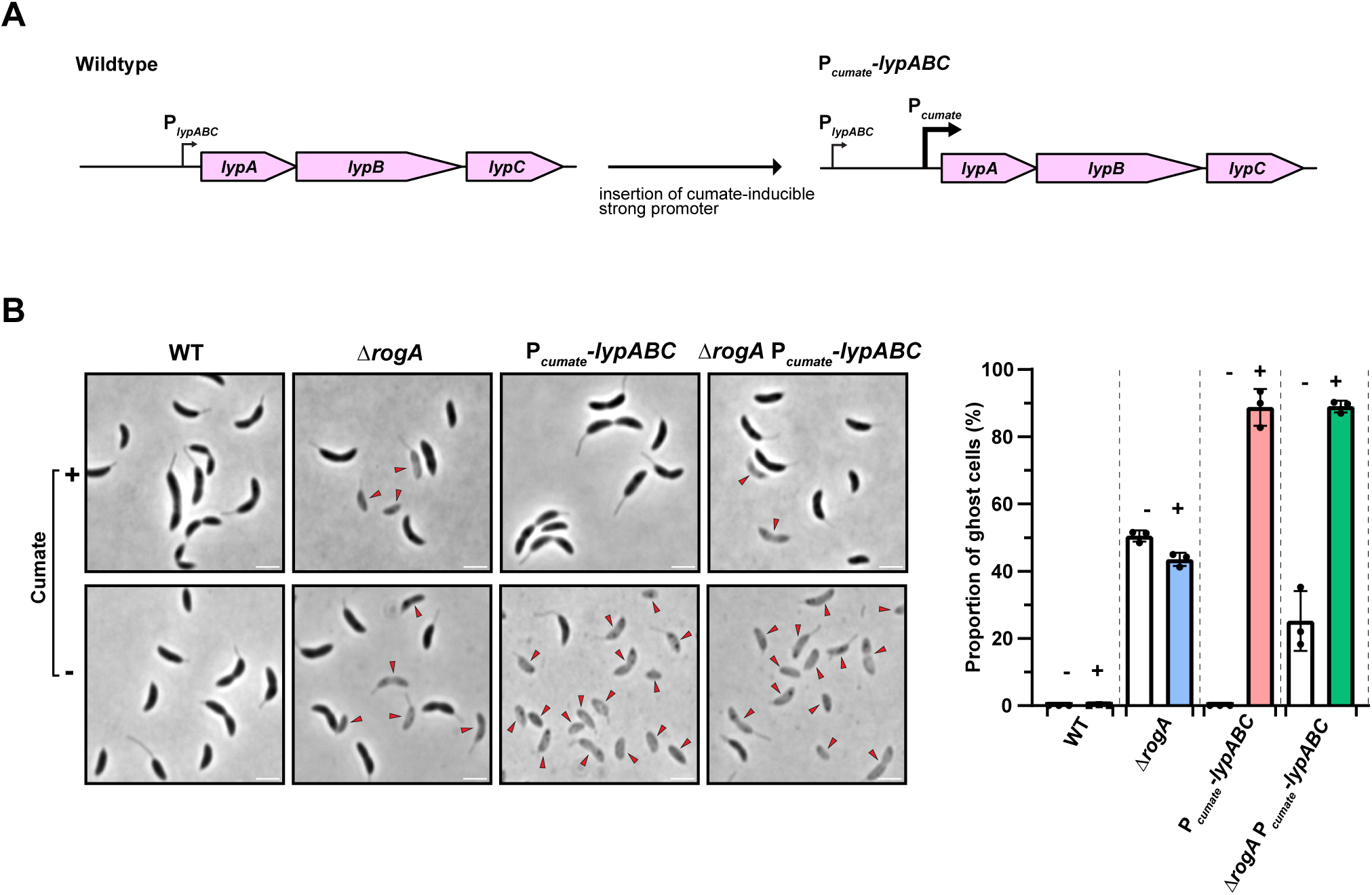
Misregulation of *lypABC* expression is highly toxic to *C. crescentus*. **(A)** Schematic to illustrate the insertion of a strong cumate-inducible promoter (P*_cumate_*) immediately upstream of the native *lypABC* locus (P*_lypABC_*). The new strain contains both the original native promoter (P*_lypABC_*) and the new P*_cumate_* promoter. **(B)** Left: *C. crescentus* strains visualised by phase-contrast microscopy. Expression of *lypABC* from the P*_cumate_* promoter was induced by the addition of 100 µM cumate to growth media. Images are representative of three independent repeats. Red arrows indicate ghost cells. Scale bar: 2 µm. Right: quantification of the proportion of ghost cells within the population of each strain. White bars: no addition of cumate. Coloured bars: addition of 100 µM cumate. Minus and plus symbols above bars also denote the absence or presence of cumate, respectively. Data represent the mean ±. s.d from three independent experiments. n=400 cells analysed per experiment.

### The XRE-like DNA-binding protein RogB represses *lypABC* expression

Given that *lypABC* overexpression results in severe cell death (**Figure 5B**), we hypothesised that *lypABC* expression must normally be tightly repressed to prevent untimely lysis in *C. crescentus*. To find potential repressors, we searched our Tn-seq dataset for genes whose disruption further reduces the fitness of Δ*rogA* cells. We identified one such candidate, CCNA_02755, encoding a predicted DNA-binding transcriptional regulator, which had fewer transposon insertions in Δ*rogA* compared to the wildtype strain (**Figure 6A**). CCNA_02755 (hereafter renamed to *rogB* for repressor <u>o</u>f <u>G</u>TA protein <u>B</u>) encodes a 132 amino acid protein containing a predicted N-terminal helix-turn-helix DNA-binding domain (aa 15-72) and C-terminal putative dimerisation domain (aa 82-132) (**Figure 6B**). RogB has since been described as an XRE-family transcriptional regulator - a protein family with multiple roles including modulation of adhesin development and susceptibility to φCbK phage infection in *C. crescentus* (*55*).

**Figure 6.**
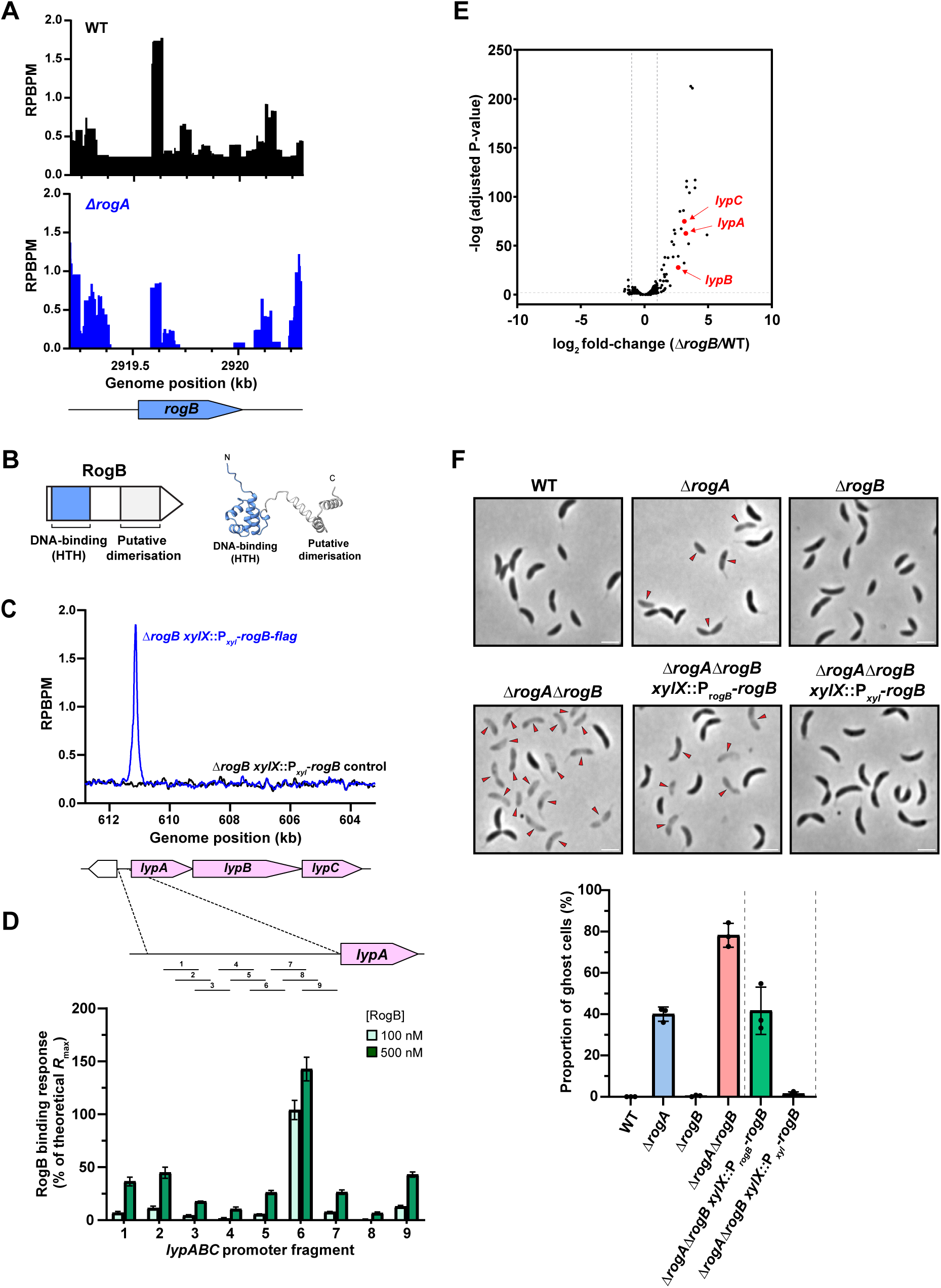
The XRE-like DNA-binding protein RogB represses *lypABC* expression. **(A)** Plot showing the reads per base pair per million reads (RPBPM) at the *rogB* genomic locus, indicating the lower frequency of transposon insertions within *rogB* in the Δ*rogA* strain (bottom, blue trace) compared to the wild-type strain (top, black trace). Three independent Tn-seq experiments were conducted. **(B)** Left: schematic of the predicted domain organisation of RogB. HTH: helix-turn-helix domain. Right: AlphaFold3 predicted structure of RogB. **(C)** ChIP-seq profiles showing RogB binding to the *lypABC* promoter *in vivo*. A *rogB-flag* (blue line) or untagged *rogB* (black line) allele was expressed from the xylose promoter (P*_xyl_*) in a Δ*rogB* mutant. Two independent repeats were performed and representative profiles are shown. **(D)** Surface plasmon resonance (SPR) experiment showing binding of purified RogB protein to the *lypABC* promoter *in vitro*. The *lypABC* promoter region was divided into nine overlapping DNA fragments. Two different concentrations of RogB protein were used: 100 nM (light green bars) and 500 nM (dark green bars). Two independent repeats were performed. **(E)** RNA-seq volcano plot showing the log_2_ fold-change for the Δ*rogB* strain compared to the wildtype strain. Grey lines indicate a value of 2.0. *lypA*, *lypB*, and *lypC*, which are upregulated when *rogB* is deleted, are annotated. Three independent repeats were performed. Top differentially expressed genes are presented as a heatmap in Figure S13A. **(F)** Top: *C. crescentus* strains visualised by phase-contrast microscopy. The Δ*rogA*Δ*rogB* deletion mutant was complemented by either expressing *rogB* from its native promoter (P*_rogB_*) or by expressing *rogB* from the xylose locus promoter (P*_xyl_*). Images are representative of three independent repeats. Red arrows indicate ghost cells. Scale bar: 2 µm. Bottom: quantification of the proportion of ghost cells within the population of each strain. Data represent the mean ±. s.d from three independent experiments. n=400 cells analysed per experiment.

To identify RogB binding sites across the genome, we performed anti-FLAG chromatin immunoprecipitation followed deep-sequencing (ChIP-seq) using a *flag*-tagged allele of *rogB* which was expressed ectopically from the *xylX* locus in a Δ*rogB* background (**Figure 6C**). Compared to an untagged RogB control, RogB-FLAG had numerous binding sites genome-wide (115 peaks with a fold-enrichment ≥2.0), including strong enrichment at the promoter of *lypABC* (**Figure 6C**, **Figure S9** and **Table S2**). To validate the ChIP-seq data, we purified RogB and tested its binding to P*_lypABC_*DNA using surface plasmon resonance (SPR). By dividing the *lypABC* promoter region into nine overlapping DNA fragments, we confirmed binding of RogB to DNA fragment 6 that was directly underneath the RogB ChIP-seq peak (**Figure 6D**). Next, to determine whether RogB activates or represses *lypABC* transcription, we performed RNA-seq comparing Δ*rogB* to wildtype cells. We observed that, while *lypABC* were transcribed at a low level in wildtype cells, *lypABC* were significantly upregulated in the Δ*rogB* mutant by 9.8-fold, 6.5-fold, and 8.6-fold, respectively, confirming that RogB acts as a transcriptional repressor of *lypABC* (**Figure 6E** and **Table S3**).

In the ghost cell-producing Δ*rogA* strain, RogB is still bound to P*_lypABC_*, thus we hypothesised that deletion of *rogB* would further activate *lypABC*, exacerbating lysis and ghost cell formation. Indeed, while *rogB* deletion alone did not produce ghost cells, the Δ*rogA*Δ*rogB* double deletion showed a significantly higher proportion of ghost cells (78.2 ± 5.8%) than the single Δ*rogA* mutant (**Figure 6F**). Complementation of Δ*rogA*Δ*rogB* with *rogB* expressed from its native promoter restored ghost cell levels to that of the Δ*rogA* mutant (41.7 ± 11.5%) (**Figure 6F**). Meanwhile, expression of *rogB* from a stronger P*_xyl_* promoter completely suppressed ghost cell formation (**Figure 6F**), likely due to further RogB-mediated repression of *lypABC*. Altogether, these findings demonstrate that RogB is a transcriptional repressor that directly binds the *lypABC* promoter region to regulate GTA-mediated host lysis.

### RogB represses *gafYZ* expression to regulate GTA activation

Further examination of the ChIP-seq data revealed that RogB also binds the promoter region of GTA activating genes *gafYZ* (P*_gafYZ_*) (**Figure 7A**). In the Δ*rogA* background, RogB binding at the *gafYZ* promoter region was further enriched by 2.6-fold, suggesting potential competitive binding between the two repressors at this promoter region (**Figure 7A** and **Table S4**). To validate the ChIP-seq data, we again performed SPR using purified RogB protein and confirmed its binding at P*_gafYZ_ in vitro* (**Figure 7B**). To determine whether RogB also represses *gafYZ* transcription, we conducted RNA-seq in a genetic background containing both a Δ*rogA* deletion to de-repress *gafYZ* and a deletion of the entire core GTA gene cluster (Δ*GTA*) to prevent the confounding loss of RNA via cell lysis. Comparing this Δ*rogA*Δ*GTA* strain to the Δ*rogB*Δ*rogA*Δ*GTA* triple mutant revealed upregulation of *gafYZ* in the absence of *rogB* by 3.5-fold and 4.0-fold, respectively (**Figure 7C** and **Table S5**). As expected and consistent with previous findings, *lypABC* were also upregulated in the Δ*rogB*Δ*rogA*Δ*GTA* strain (**Figure 7C**). Finally, immunoblotting with an anti-GafY polyclonal antibody (*36*) confirmed that GafY protein was more abundant in the absence of *rogB* (**Figure 7D**). Overall, our data demonstrate that RogB directly binds to and represses the transcription of GTA activating genes *gafYZ*. RogB therefore acts as a dual repressor, coupling two different GTA life stages: transcriptional activation via GafYZ and host cell lysis via LypABC.

**Figure 7.**
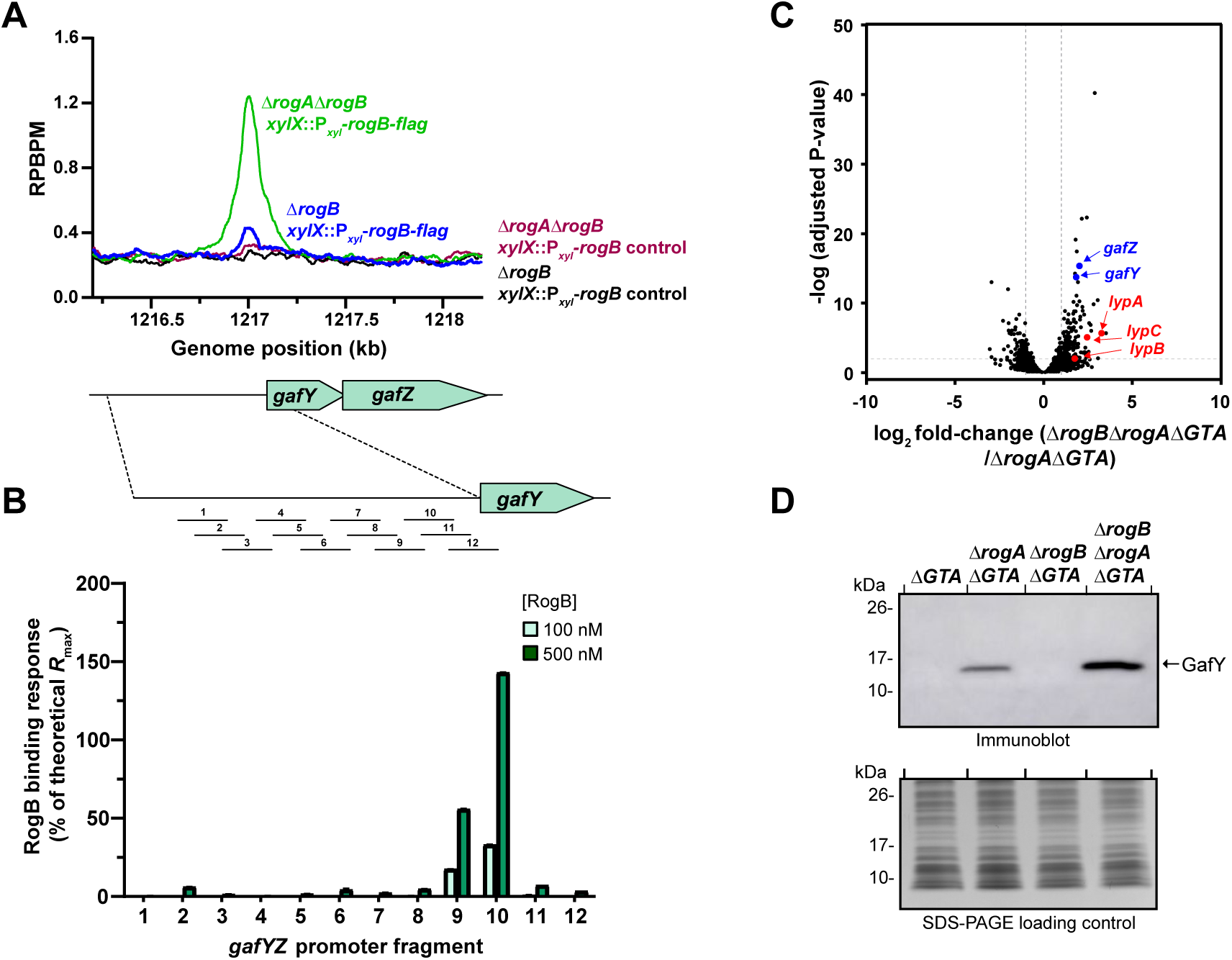
RogB represses *gafYZ* expression to regulate GTA activation. **(A)** ChIP-seq profiles showing binding of RogB to the *gafYZ* promoter *in vivo*. A *rogB-flag* allele was expressed from the xylose promoter (P*_xyl_*) in either a single Δ*rogB* mutant (blue line) or a double Δ*rogA*Δ*rogB* mutant (green line). Untagged *rogB* controls are shown in black and maroon for the single and double mutant backgrounds, respectively.Two independent repeats were performed and representative profiles are shown. **(B)** Surface plasmon resonance (SPR) experiment showing binding of purified RogB protein to the *gafYZ* promoter *in vitro*. The *gafYZ* promoter region was divided into twelve overlapping DNA fragments. Two different concentrations of RogB protein were used: 100 nM (light green bars) and 500 nM (dark green bars). Two independent repeats were performed. **(C)** RNA-seq volcano plot showing the log_2_ fold-change for the Δ*rogB*Δ*rogA*Δ*GTA* strain compared to the Δ*rogA*Δ*GTA* strain. Grey lines indicate a value of 2.0. *lypA*, *lypB*, *lypC*, *gafY*, and *gafZ* which are upregulated when *rogB* is deleted, are annotated. Three independent repeats were performed. Top differentially expressed genes are presented as a heatmap in Figure S13B. **(D)** Immunoblot (top) of total cell lysates from *C. crescentus* using a polyclonal anti-GafY antibody to show the levels of GafY protein in the indicated mutants. A separate Coomassie-stained SDS-PAGE gel was loaded with equal sample volumes to serve as a loading control (bottom). The immunoblot is representative of at least two independent experiments.

## DISCUSSION

### A new mechanism of bacterial cell lysis for the release of GTA particles

Our work uncovers a mechanism of bacterial cell death in which domains typically associated with anti-phage immunity appear to have been co-opted to instead facilitate the release of GTA particles. Central to this mechanism is the LypABC system which is tightly regulated by a transcriptional repressor, RogB, to ensure controlled lysis. Catalytic activity of each LypABC domain, excepting the endonuclease activity of LypC, is essential for lysis, suggestive of a complex multipartite pathway. Notably, the rarity of LypABC homologues in other GTA-producing bacteria suggests that LypABC-mediated host cell lysis may not be universal and that alternative lysis pathways likely exist. Indeed, in the *Rhodobacter capsulatus* GTA system, host cell lysis occurs via a phage-like holin-endolysin mechanism (*23*).

The precise mechanism by which LypABC - and in fact most CARD-NLR systems - are activated and lead to cell death is currently unknown. Since LypABC are produced in GTA-off wildtype cells, we reason that the system remains autoinhibited until GTA activation occurs to prevent untimely and unproductive host cell lysis (**Figure 8A**). Since many anti-phage defence systems are often activated by direct interaction with phage structural proteins (*48, 56–60*), it is possible that a similar mechanism involving GTA structural components might underlie LypABC activation, however extensive co-immunoprecipitation with mass spectrometry and bacterial two-hybrid screens have thus far failed to identify such direct protein activators. Sensing of phage infection can also occur indirectly through defence system guarding of host proteins and processes (*61–63*), therefore it is possible that LypABC activation occurs indirectly via sensing of host cell perturbations such as GTA-specific host DNA damage or other stress signals.

**Figure 8.**
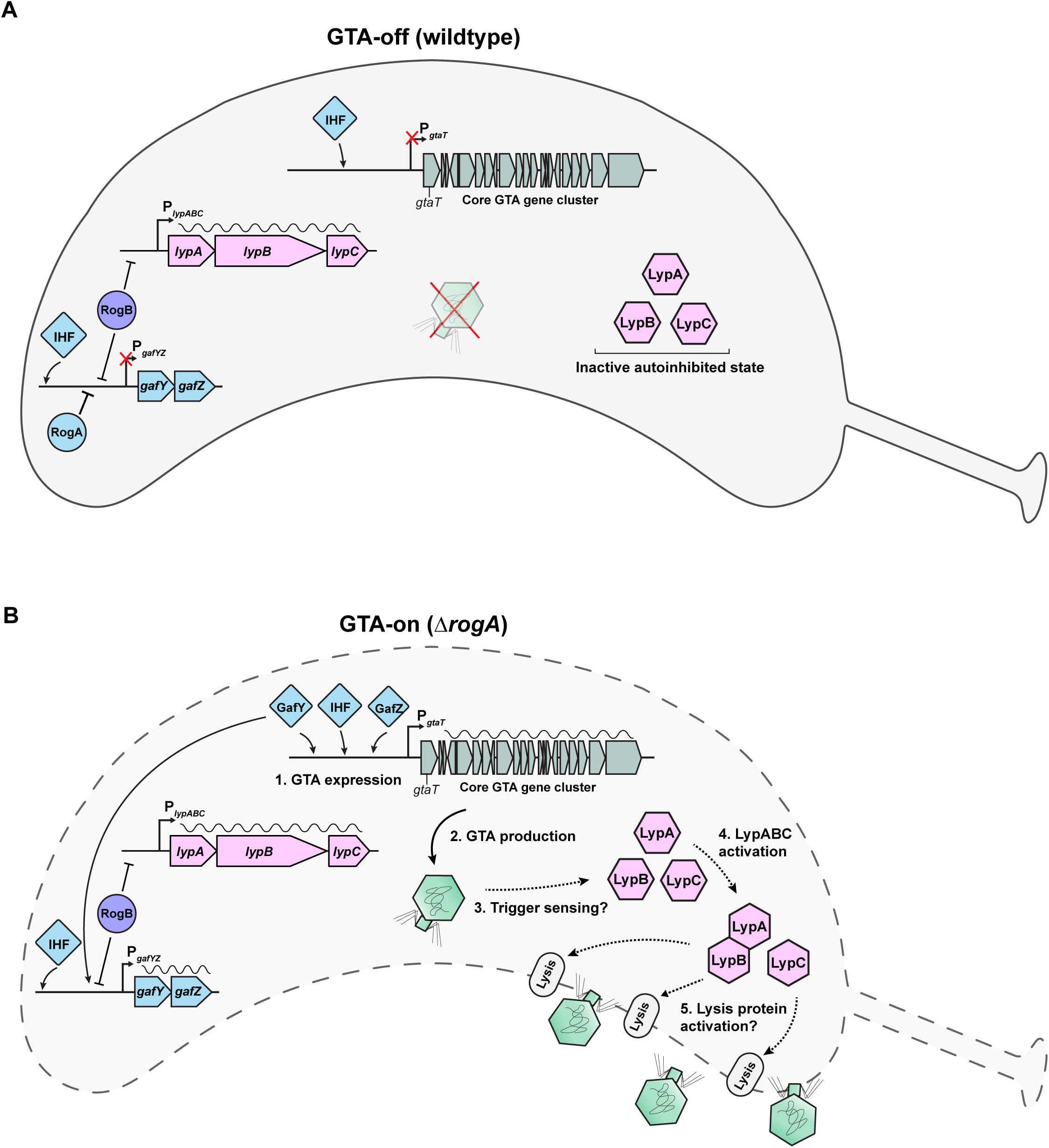
A proposed model for GTA production and host cell lysis. Diagram to illustrate our current model of how GTA expression is regulated and how LypABC function to facilitate host cell lysis. **(A)** GTA-off wildtype strain. Repressor proteins RogA and RogB bind to and repress transcription from the *gafYZ* promoter (P*_gafYZ_*). IHF binding to P*_gafYZ_* and to the core GTA cluster promoter (P*_gtaT_*) is insufficient to activate gene transcription from either operon. Basal transcription of *lypABC* occurs, despite binding of RogB to the *lypABC* promoter (P*_lypABC_*). LypABC proteins are produced but most likely in an inactive autoinhibited state. GTAs are not produced and cells do not lyse. **(B)** GTA-on*ΔrogA* strain. Transcriptional repression of *gafYZ* is relieved due to the deletion of the *rogA* repressor. GafYZ bind to and activate transcription of the core GTA gene cluster (1). Basal transcription of *lypABC* occurs, despite binding of RogB to the *lypABC* promoter (P*_lypABC_*). GTA particles are produced (2) and may directly or indirectly trigger (3) the activation of LypABC proteins (4). LypABC may then activate lysis protein(s) that allow the release of GTA particles into the extracellular environment (5). Black wavy lines above gene operons represent mRNA transcription. Black solid or dashed arrows: steps for which a mechanism is known or unknown, respectively.

Overexpression of *lypABC* caused severe levels of cell lysis including in non-GTA-producing wildtype cells (**Figure 5**). In plants, overexpression of NLR proteins is known to shift NLRs from an autoinhibited state to an active state, causing a hypersensitive response that leads to cell death even in the absence of pathogens (*64–66*). It is therefore possible that excess LypABC may mirror this shift to an active state, initiating *C. crescentus* cell death autonomously.

In CARD-NLR anti-phage systems, cell death is directly caused by death-domain effectors such as gasdermins, phospholipases, or endonucleases (*29, 44*). As the LypABC system does not contain a gasdermin protein and LypC endonuclease activity is dispensable, LypABC proteins do not appear to directly execute cell lysis. Instead, we propose a model in which LypABC indirectly sense GTA production, transitioning from a resting autoinhibited state to an active state (**Figure 8B**). Once activated, LypABC likely trigger the activity of downstream lytic effector(s) (whose identity is current unknown), resulting in cell lysis. Work is ongoing to elucidate the mechanisms underpinning how LypABC senses GTA production and initiates the downstream lytic cascade.

### Multi-layered regulation of GTA life stages

Host cell lysis is not the only GTA life stage under tight regulatory control; GTA activation in *C. crescentus* is completely inhibited under laboratory conditions via RogA-mediated repression of *gafYZ* expression (*26*). Although the natural environmental stimuli that trigger GTA activation remain elusive, our study here identifies another regulatory layer wherein RogB binding to the *gafYZ* promoter further modulates GTA expression.

Deletion of *rogB* alone is insufficient for GTA activation (**Figure 6**), indicating that RNA polymerase is unable to overcome RogA-mediated repression. However, deletion of both *rogA* and *rogB* together resulted in significantly more ghost cells, suggesting that RogB reinforces *gafYZ* repression. Given that not all cells lyse in the Δ*rogA*Δ*rogB* double mutant, it is likely that additional repressors control the key *gafYZ* promoter. Indeed, prior work identified additional XRE-family proteins such as RtrA, RtrB, and CdxA, which can repress *gafYZ* transcription when overexpressed (*55*). The XRE-family, which includes RogB, has multiple binding sites genome-wide (*55*). It is therefore possible that these repressors - which also have other roles in *C. crescentus* physiology - may have been co-opted to refine the transcriptional regulation of GTA activation. Our findings support a model in which RogB functions as a dual repressor of both GTA activation (via *gafYZ*) and GTA-mediated lysis (via *lypABC*), ensuring that the two critical processes are coupled (**Figure 8**).

### Repurposing of viruses and immune systems

Domestication of phages resulted in GTAs, a striking example of viruses that have been de-weaponised to benefit bacterial hosts. In nature, viral domestication is not uncommon; phage tails have been repurposed into tools for bacterial warfare such as type VI secretion systems (*67*) and tailocins (*68*). Other examples include co-option of a phage protein to benefit bacterial endosymbionts of marine sponges by aiding immune evasion (*69*), or detoxification of a phage lysin that resulted in a new bacterial morphogenesis protein (*70*). Humans also contain domesticated endogenous viral elements (*71*) such as the retroviral envelope protein, syncytin, which has been co-opted for placental development (*72*). In eukaryotes, immune pathways are very versatile systems, having been repeatedly repurposed for alternative functions beyond host defence (*73, 74*). For example, neuronal major histocompatibility complex (MHC) class I immunity molecules are employed for the development and plasticity of the central nervous system (*75, 76*). Moreover, nucleotide-binding and leucine-rich repeat (NLR)-based innate immunity proteins also function in non-immune roles such as transcriptional regulation and reproduction (*77–80*).

LypABC-mediated cell lysis appears to share some similarity with anti-viral immune systems that confer overall population immunity via death of infected individual cells. While we cannot fully exclude a role for LypABC in anti-phage defence, our evidence strongly supports a role for LypABC in facilitating the release of GTA particles. Overall, our work highlights the plasticity of immune systems, demonstrating that bacterial immune components can evolve new roles as positive drivers of MGEs.

## MATERIALS AND METHODS

### Strains, media, and growth conditions

*Escherichia coli* was routinely grown in LB media at 37 °C and *Caulobacter crescentus* strain NA1000 was grown in PYE media at 30 °C. To culture *C. crescentus* cells to stationary phase for GTA production, 10 ml of PYE liquid was inoculated with a single colony and grown at 30 °C for 20 h with shaking at 250 rpm. A small volume (calculated to set a starting OD_600_ of 0.1) was then transferred from this initial culture into 10 ml of fresh PYE media and grown for a further 20 h. If required, *C. crescentus* growth media were supplemented with antibiotics at the following concentrations (liquid/plates): kanamycin 5/25 μg/ml, oxytetracycline 1/1 μg/ml, spectinomycin 25/100 μg/ml, and carbenicillin n.a./50 μg/ml. Concentrations used for *E. coli* growth media (liquid/plates) were: kanamycin 50/50 μg/ml, spectinomycin 50/50 μg/ml, chloramphenicol 20/30 μg/ml, and carbenicillin 50/100 μg/ml. To induce or repress gene expression from the *C. crescentus* P*_xyl_* promoter, liquid PYE media were supplemented with either 0.3% xylose (final concentration) or 0.2% glucose (final concentration), respectively. To induce gene expression from the *C. crescentus* P*_cumate_*promoter, 100 μM cumate (final concentration) was added to liquid PYE media. Transformations and transductions into *C. crescentus* were performed as previously described (*81*). All strains, plasmids and oligonucleotides that were constructed and used are listed in **Table S6, Table S7** and **Table S8**, respectively. Complete details of plasmid and strain construction are available in the Supplementary Information.

### Genomic DNA extraction

To extract total genomic DNA from *C. crescentus* strains, 1 ml of stationary phase cells was centrifuged at 17,000 x *g* for 1 min and the pellet was resuspended in 300 μl of Cell Lysis Solution (PureGene, Qiagen). Resuspended cells were incubated at 50 °C for 15 min, mixed with 1 μl of RNase A (NEB 20 mg/ml stock) and incubated at 37 °C for 1 h. Samples were cooled to room temperature, mixed with 200 μl of Protein Precipitation Solution (PureGene, Qiagen) and then centrifuged at 17,000 x *g* for 10 min. The supernatant was combined with 600 μl of isopropanol and mixed well by inversion. Samples were centrifuged again at 17,000 x *g* for 1 min and the supernatant discarded. Pellets were resuspended in 600 μl of 70% ethanol and centrifuged at 17,000 x *g* for 1 min. The supernatant was discarded, and a final 1 min centrifugation was performed to remove any remaining liquid. The DNA pellet was resuspended in 100 μl of sterile nuclease-free water and further incubated at 37 °C for 15-30 min. To assay for encapsulation of host DNA into GTA particles, 50 μl of genomic DNA was run on a 1% agarose gel at 150 V for 45-60 min and visualised to determine the presence/absence of an 8.3 kb band.

### Transposon mutagenesis with deep sequencing (Tn-seq)

To prepare strains for Tn-seq, 10 ml of *C. crescentus* NA1000 wildtype and Δ*rogA* strains were cultured to stationary phase; simultaneously the Tn5 transposon delivery plasmid pMCS-6-Tn5-R6kg-kan2 was introduced into an *E. coli* S17-1 conjugative donor strain by heat-shock transformation. The next day, *E. coli* transformants were resuspended in LB liquid and washed three times by centrifugation to remove kanamycin. *C. crescentus* and *E. coli* cultures were adjusted to an OD_600_ of 0.5 and then 10 ml of each *C. crescentus* strain was mixed with 1 ml of *E. coli* S17-1 harbouring pMCS-6-Tn5-R6kg-kan2. The mixtures were centrifuged at 4,650 x *g* for 5 min and the pellet was resuspended in 500 μl of PYE. Ten sterile 0.45 μm nitrocellulose membranes were placed onto PYE agar plates and then 50 μl of the *C. crescentus/E. coli* resuspension was pipetted onto each membrane and incubated at 30 °C for 4 h. Cells were removed from each filter membrane by the addition of 500 μl of PYE followed by vigorous vortexing and then combined. Five hundred microlitres of either the wildtype or Δ*rogA* conjugation mix were spread onto large PYE agar plates supplemented with kanamycin (to select for *C. crescentus* cells with transposon insertions) and carbenicillin (to kill the *E. coli* S17-1 donor) and incubated at 30 °C for 3 days. Resulting colonies were washed from large Petri disks and resuspended in PYE liquid, mixed thoroughly, and 1 ml removed for total genomic DNA extraction. Purified genomic DNA was sheared into 200-500 bp fragments and libraries were constructed for Illumina sequencing as previously described (*82*) followed by a PCR amplification using a universal P5-ME primer annealing to the transposon and a specific index primer annealing to adaptor-ligated sheared DNA fragments. DNA was sequenced on an Illumina HiSeq2500 platform at the Tufts University Genomics facility. To analyse Tn-seq data, short single-end Illumina reads were mapped back to the *C. crescentus* NA1000 reference genome using HISAT2 (*83*) and then associated with genes using HTSEQ-count. Log_2_ fold-change and P-adjusted values were calculated using DESeq2 (*84*) and a custom Python script was used to annotate genes in the DESeq2 output file.

### RNA sequencing (RNA-seq)

*C. crescentus* strains were cultured to stationary phase and 5 ml cell cultures were pelleted for RNA extraction. Total RNA was purified using the Direct-zol RNA miniprep kit (Zymo Research) and 10 μg was incubated with 20 units of Turbo DNaseI (Invitrogen) at 37 °C for 1 h to remove any contaminated genomic DNA. DNaseI was subsequently removed using the RNA clean and concentrator-25 kit (Zymo Research). Purified RNA samples were shipped to Genewiz where bacterial rRNA depletion was performed with an NEBNext rRNA Depletion Kit (Cat. No. E7850X). DNA libraries were prepared by Genewiz and sequenced on an Illumina NovaSeq platform. RNA-seq data, consisting of short, paired-end Illumina reads, were analysed as described previously for the Tn-seq data.

### Chromatin immunoprecipitation with deep sequencing (ChIP-seq)

*C. crescentus* strains were cultured to stationary phase and fixed with a final concentration of 1% formaldehyde. Cells were incubated on a wheel rotator at room temperature for 30 min, then the reaction was quenched by the addition of 250 mM glycine for 10 min. Cells were washed three times in 1x PBS and then resuspended in 1 ml of cell lysis buffer 1 (20 mM K-HEPES pH 7.9, 50 mM KCl, 10% glycerol, and Roche EDTA-free protease inhibitors). Samples were sonicated on ice (11 cycles of 15 s “on” and 15 s “off” at an amplitude setting of 8) using a Soniprep 150 probe-type sonicator, then debris was pelleted by centrifugation at 17,000 x *g* at 4 °C for 20 min. Supernatant was transferred to a new microcentrifuge tube and the buffer concentration was adjusted via the addition of: 10 μl of 1 M Tris pH 8 (10 mM final concentration), 20 μl of 5 M NaCl (150 mM final concentration), and 10 μl of 10% NP40 (0.1% final concentration). Fifty microlitres were then transferred to a new tube (“input” control) and stored at −20 °C. Storage buffer was removed from α-FLAG antibody M2 agarose beads (Merk) by repeated cycles of centrifugation and resuspension of α-FLAG beads (100 μl) in 1 ml of IPP150 buffer (10 mM Tris-HCl pH 8, 150 mM NaCl, and 0.1% NP40). ChIP sample supernatant was transferred to the prepared α-FLAG beads and incubated on a wheel rotator at 4 °C overnight. Samples were then washed five times with 1 ml of IPP150 buffer, followed by two washes with 1 ml of 1x TE buffer (10 mM Tris-HCl pH 8, 1 mM EDTA). A two-step elution of protein-DNA complexes was performed. Briefly, beads were first incubated with 150 μl of elution buffer (50 mM Tris-HCl pH 8, 10 mM EDTA, and 1% SDS) at 65 °C for 15 min, then centrifuged at 17,000 x *g* for 5 min, and the supernatant transferred to a new tube. Beads were then incubated with 100 μl of 1x TE + 1% SDS at 65°C for an additional 15 min, then centrifuged at 17,000 x *g* for 5 min. Supernatant eluates (ChIP samples) were combined and incubated at 65 °C overnight to reverse crosslinks. The “input” control fraction was also incubated at 65 °C overnight with the addition of 200 μl of 1x TE + 1% SDS. DNA from the ChIP and input fractions was purified with a QIAquick PCR purification kit (Qiagen) and eluted in 40 μl of water. Libraries were then prepared for Illumina sequencing using the NEXT Ultra II library preparation kit (NEB). DNA was sequenced on an Illumina Nextseq 550 platform at the Tufts University Genomics facility. To analyse ChIP-seq data, short paired-end Illumina reads were mapped back to the *C. crescentus* NA1000 reference genome using HISAT2. The sequencing coverage for each nucleotide position was determined using bedtools genome-cov (*85*). Peaks were identified using MACS2 callpeak (*86*). ChIP-seq profiles were plotted using GraphPad Prism, with the x-axis representing the genomic nucleotide position and the y-axis representing the reads per base pair per million reads (RPBPM).

### Protein purification

Plasmid pET21b::*rogB-his6* was introduced into *E. coli* Rosetta BL21 DE3 (pLys) cells by heat-shock transformation. A 20 ml overnight culture was used to inoculate 1 l of LB media supplemented with chloramphenicol and carbenicillin. Cells were cultured at 37 °C with shaking to early exponential phase (OD_600_ = ∼0.4), cooled to 18 °C, and then supplemented with 1 mM isopropyl-B-D-thiogalactopyranoside (IPTG). The culture was then incubated at 18 °C with shaking for 20 h to induce protein production. Cells were harvested by centrifugation at 4,248 x *g* for 10 min and resuspended in 25 ml of buffer A (5% glycerol, 100 mM Tris-HCl pH 8.0, 300 mM NaCl, 10 mM imidazole) supplemented with one EDTA-free protease inhibitor tablet and lysozyme. Cells were then lysed by sonication (10 cycles of 15 s, resting on ice for 15 s between cycles). Cell debris was pelleted by centrifugation at 32,000 x *g* for 35 min and the supernatant filtered through a 0.22 μm membrane. The lysate was incubated with 2 ml of HIS-Select Cobalt Affinity Gel resin (Merck) (pre-washed with 50 ml of buffer A) on a wheel rotator at 4 °C for 1 h. The lysate was then drained and the resin washed with 75 ml of buffer A. Protein was eluted in 2.7 ml of buffer B (5% glycerol, 100 mM Tris-HCl pH 8.0, 300 mM NaCl, 500 mM imidazole). Finally, protein was de-salted using a PD-10 column, checked for purity by SDS-PAGE, and stored at −80 °C in storage buffer (5% glycerol, 100 mM Tris-HCl pH 8.0, 300 mM NaCl).

### Surface plasmon resonance (SPR)

SPR experiments were performed using a Biacore 8K (Cytiva) instrument using the previously-developed Reusable DNA Capture Technique (ReDCaT) (*87*) to detect protein-DNA interactions. Briefly, the promoter regions of *gafYZ* and *lypABC* were divided into 50 bp single-stranded DNA oligonucleotides that overlap by 20 bp. These oligonucleotides were annealed to a complementary ssDNA fragment containing a ReDCaT-specific adaptor extension (*87*). The adaptor binds to a biotinylated linker which is permanently bound to the SPR chip surface, enabling each DNA fragment to bind and be stripped off the chip after each experiment. Double-stranded DNA duplexes were diluted to 1 μM in 1x HBS-EP+ buffer (150 mM NaCl, 3 mM EDTA, 0.005% Tween-20, 10 mM HEPES, pH 7.4) and bound to the SPR chip. Purified RogB protein was diluted in 1x HBS-EP+ buffer to either 100 nM or 500 nM final concentration and then flowed over the DNA-bound chip, followed by a 1x HPS-EP+ buffer-only wash to allow protein dissociation. Finally, both protein and DNA were removed using a regeneration solution (1 M NaCl and 50 mM NaOH) wash to regenerate the chip. Binding events were recorded in response units and the strength of protein-DNA binding was expressed as a percentage of the theoretical *R*_max_ (*87*).

### Immunoblots

Bacterial cells were harvested by centrifugation at 17,000 x *g* for 10 min and pellets resuspended in 300 μl of lysis buffer (20 mM K-HEPES pH 7.9, 50 mM KCl, 10% glycerol, one EDTA-free protease inhibitor tablet). Cells were lysed by sonication (3 cycles of 10 s, resting on ice for 10 s between cycles) and debris pelleted by centrifugation at 17,000 x *g* for 15 min at 4 °C. If required, extracellular fractions were prepared by filtering supernatant from harvested cultures through a 0.22 μm membrane, following by spin-concentration in Amicon Ultra-15 centrifugal filter units with a 100 kDa molecular weight cut-off to collect GTA particles. Total protein concentrations were determined using Bradford reagent and used to match the amount of total protein loaded across different samples. Samples were denatured by boiling in SDS-PAGE loading dye containing β-mercaptoethanol and then loaded onto 12% Novex Tris-Glycine WedgeWell gels (ThermoFisher) alongside a Color Pre-stained Broad Range protein ladder (NEB). Gels were run at 200 V for 30 min, then proteins were transferred onto a PVDF membrane using the Trans-Blot Transfer System (BioRad). The membrane was blocked in 1x TBS buffer (100 mM Tris-HCl, 140 mM NaCl, pH 7.4) + 0.1% Tween-20 + 5% milk powder) for 1 h. For α-FLAG immunoblots, the membrane was then incubated with a 1:5000 dilution of a monoclonal α-FLAG M2-Peroxidase HRP-conjugated antibody (Merck) for 1 h. The membrane was then washed five times in TBS buffer + 0.1% Tween-20) for 1 min each time. Finally, the membrane was incubated with SuperSignal West Femto Maximum Sensitivity Substrate (Thermo Scientific) and visualised in an Amersham Imager 600 (GE Healthcare). For immunoblots to detect either GafY or GtaL, the membrane was first incubated with either a custom-made polyclonal α-GafY antiserum (3:1000 dilution) or polyclonal α-GtaL antiserum (1:1000 dilution) for 1 h. Following three 5 min washes in TBS buffer + 0.1% Tween-20, membranes were incubated with a 1:10000 dilution of HRP-conjugated goat anti-rabbit antibody (cat# ab6721, Abcam) for 1 h. Membranes were washed three times for 5 min and imaged as described above. Protein loading controls were run on 12% SDS-PAGE gels and stained with an InstantBlue Coomassie protein stain (Abcam).

### Light microscopy and image analysis

Bacteria were immobilised on 1% agarose pads and visualised under a Zeiss Axio Observer Z.1 inverted epifluorescence microscope equipped with a Zeiss Plan Apochromat 100x/NA 1.4 Ph3 objective lens, an sCMOS camera (Hamamatsu Orca FLASH 4), and a Zeiss Colibri 7 LED light source. For PI staining, cells were incubated with a final concentration of 15 μM PI for 15 min in the dark and then immediately imaged on agarose pads. The following filter sets were used when required: GFP (excitation: 450-488 nm, emission 499-549 nm) and PI (excitation: 450-488 nm, emission 599-659 nm). Images were acquired in Zeiss Zen Blue software then processed and analysed in Fiji (*88*). The MicrobeJ plug-in (*89*) for Fiji was used for quantitative analysis. All bacteria detected in MicrobeJ (n=400 per biological repeat) were inspected to ensure that they were detected correctly. For ghost cell quantification, the mean intensity value of the phase channel was automatically measured in each individual cell and a threshold was set to distinguish between phase-light ghost cells (lower intensity value) and phase-dark cells (higher intensity value). To determine the proportion of ghost cells, the number of cells with a mean phase intensity value beneath the threshold was calculated and expressed as a percentage of the total cell population. To quantify the proportion of mNG-fluorescent or PI-fluorescent cells, the mean intensity value of each fluorescence channel was automatically measured in each individual cell and a threshold was set to distinguish between fluorescent and non-fluorescent cells. To determine the proportion of fluorescent cells, the number of cells with a mean fluorescence intensity value above the threshold was calculated and expressed as a percentage of the total cell population.

### Time-lapse microscopy

To prepare strains for time-lapse microscopy, a 10 ml culture of *C. crescentus* was grown for 16 h at 30 °C with shaking. A 5 ml aliquot from this culture was then centrifuged at 4,650 x *g* for 10 min and the supernatant filtered through a 0.22 μm membrane to collect early stationary-phase spent PYE culture media. Spent PYE media was then mixed with melted agarose and pipetted into a GeneFrame (Thermo Scientific) attached to a glass slide to generate a 1.2% PYE agarose pad. Where required, the agarose pad was supplemented with PI at a final concentration of 0.1 μM. Once set, 1 μl of *C. crescentus* culture was pipetted onto the agarose pad which was then covered with a cover slip and firmly sealed. Time-lapse experiments were performed at 30 °C inside a temperature-controlled incubation chamber fitted to the Zeiss Axio Observer Z.1 microscope. Time-lapse images were acquired every 10 min across 10 different positions. Data were acquired in Zeiss Zen Blue software then processed and analysed in Fiji. Where required, channels were registered to correct for drift using the HyperStackReg plug-in for Fiji.

### Cryo-electron tomography and processing

*C. crescentus* strains were grown for 20 h, diluted in fresh media to an OD_600_ of 0.1 and then incubated for a further 20 h. Cultures were mixed in a ratio of 1:2 with 6 nm BSA-conjugated gold fiducials (Aurion), which were buffer transferred to PYE. The mixture (3.8 μl) was applied to 200-mesh R2/1 copper grids (Quantifoil) and vitrified in liquid ethane (blot time of 7 s, force 0, wait time of 2 s) using a Vitrobot Mark IV (Thermo Fisher Scientific). The sample was blotted using only the back vitrobot pad, with the front pad covered in parafilm as reported previously (*90*). Grids were either imaged at 200 kV using a Glacios TEM equipped with a Falcon4 detector (University of York) or imaged at 300 kV using a Krios G3i TEM equipped with a K3 camera (SLAC-Stanford). Tilt series were acquired using a dose symmetric scheme (SLAC-Stanford data) or bidirectional scheme starting from −21° (University of York data), with 3° increment and 54°/-54° span. The dose per tilt image was 2.9 (SLAC-Stanford data) and 3.2 e^-^Å^-2^ (University of York data). Tilt images were gain corrected, aligned to stacks and dose weighted using the IMOD alignframes command (*91*). Stacks were imported to EMAN2 (*92*) where tomograms were reconstructed. Tomograms were visualised using IMOD. Tomogram segmentation was performed in EMAN2 using the tomoseg convolutional neural network-based semi-automated cellular tomogram annotation protocol (*93*) and segmented volumes were visualised using ChimeraX (*94*).

### Bacterial two-hybrid assay

Bacterial two-hybrid assays were performed exactly as described in the Euromedex Bacterial Adenylate Cyclase Two-Hybrid System Kit Manual (Cat No: EUK001). Briefly, *E. coli* BTH101 cells were co-transformed with a pair of plasmids by electroporation, then spread onto LB plates containing carbenicillin and kanamycin and incubated at 30 °C for 20 h. Three colonies from each plasmid combination were cultured in LB with carbenicillin and kanamycin for 16 h, then 5 μl were spotted on McConkey agar plates and incubated at 30 °C for 24 h.

### Bioinformatics analysis and structural predictions

Sequence similarity searches for LypABC protein homologues were carried out in BLASTP with default parameters and structural homology searches were performed with Foldseek (*45*). The tools InterProScan (*95*) and HHPred (*96*) were used to annotate protein domains. AlphaFold3 (*97*) was used to generate predicted protein structures. Models were visualised in PyMOL v.2.5.3 (Schrödinger) and prepared for presentation using UCSF ChimeraX v.1.9 (*94*).

## ACKNOWLEDGMENTS

We thank members of our laboratory, Michael Laub, Mark Buttner, Sophien Kamoun, Jonathan Jones, and Susan Schlimpert for helpful discussion and feedback on this manuscript. We thank Rebecca Lo for preliminary strain construction, and Jovana Kaljevic and Boris Stojilkovic for guidance with RNA-seq analysis. This work is supported by a Lister Institute Fellowship and Wellcome Trust Investigator grant 221776/Z/2/Z (to T.B.K.L.), a Royal Commission for the Exhibition of 1851 Fellowship (to E.J.B), a Sir Henry Wellcome Fellowship (224067/Z/21/Z to P.B), and the BBSRC-funded Harnessing Biosynthesis for Sustainable Food and Health (HBio) Institute Strategic Programme BB/X01097X/1 (to the John Innes Centre). The cryo-EM imaging at the University of York cryo-EM facility was supported by the Wellcome Trust (206161/Z/17/Z); and at Stanford-SLAC Cryo-EM Center (S2C2) by the National Institute of General Medical Sciences (1R24GM154186). The content is solely the responsibility of the authors and does not necessarily represent the official views of the National Institutes of Health. The authors would also like to thank the following S2C2 personnel for their invaluable support and assistance: Dr Yan Liu, Prof Wah Chiu.

## Data and materials availability

The ChIP-seq, RNA-seq and Tn-seq raw data generated in this work have been deposited in the GEO database under the accession codes GSE295577, GSE295580, and GSE295581, respectively. All plasmids and strains constructed in this study are available upon request.

## Author contributions

Conceptualization: EJB, TBKL

Methodology: EJB, PB, NTT, PMN, AM, TBKL

Investigation: EJB, PB, NTT, PMN, AM, TBKL Visualization: EJB, PB, PMN

Funding acquisition: EJB, PB, TBKL Project administration: EJB, TBKL Supervision: TBKL

Writing – original draft: EJB, PB, NTT, PMN, AM, TBKL Writing – review & editing: EJB, TBKL

---

**Figure S1.**
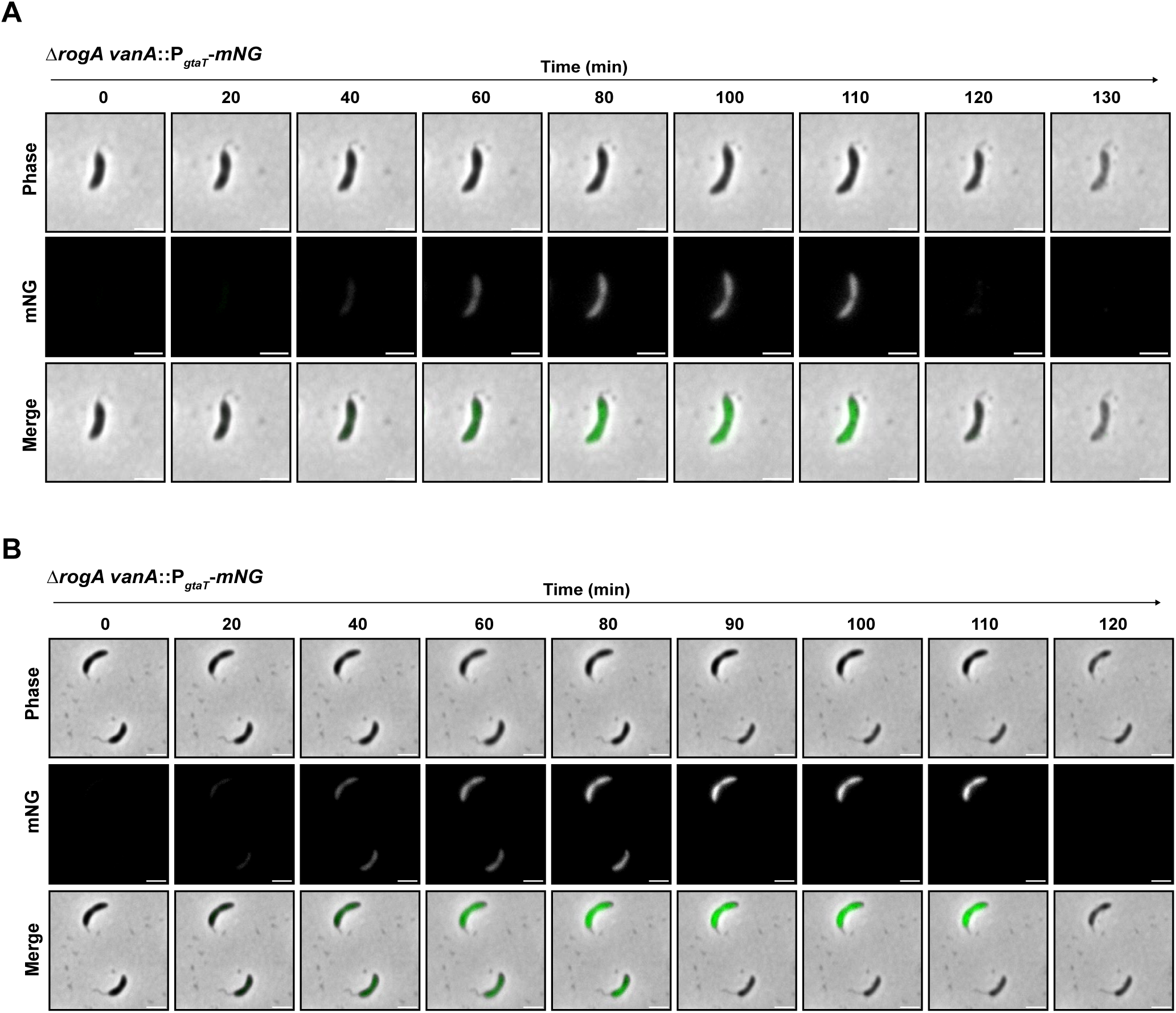
GTA production and host cell lysis in *Caulobacter crescentus* cells. Two additional examples of fluorescence microscopy images from time-lapse experiments performed with the Δ*rogA vanA*::P*_gtaT_* -*mNG* strain showing GTA activation (mNG signal) followed by cell lysis (transition to phase-light ghost cell). Time-lapses can be viewed in Movie S2 **(A)** and Movie S3 **(B)**. Data are representative of at least three independent repeats. Scale bar: 2 µm.

**Figure S2.**
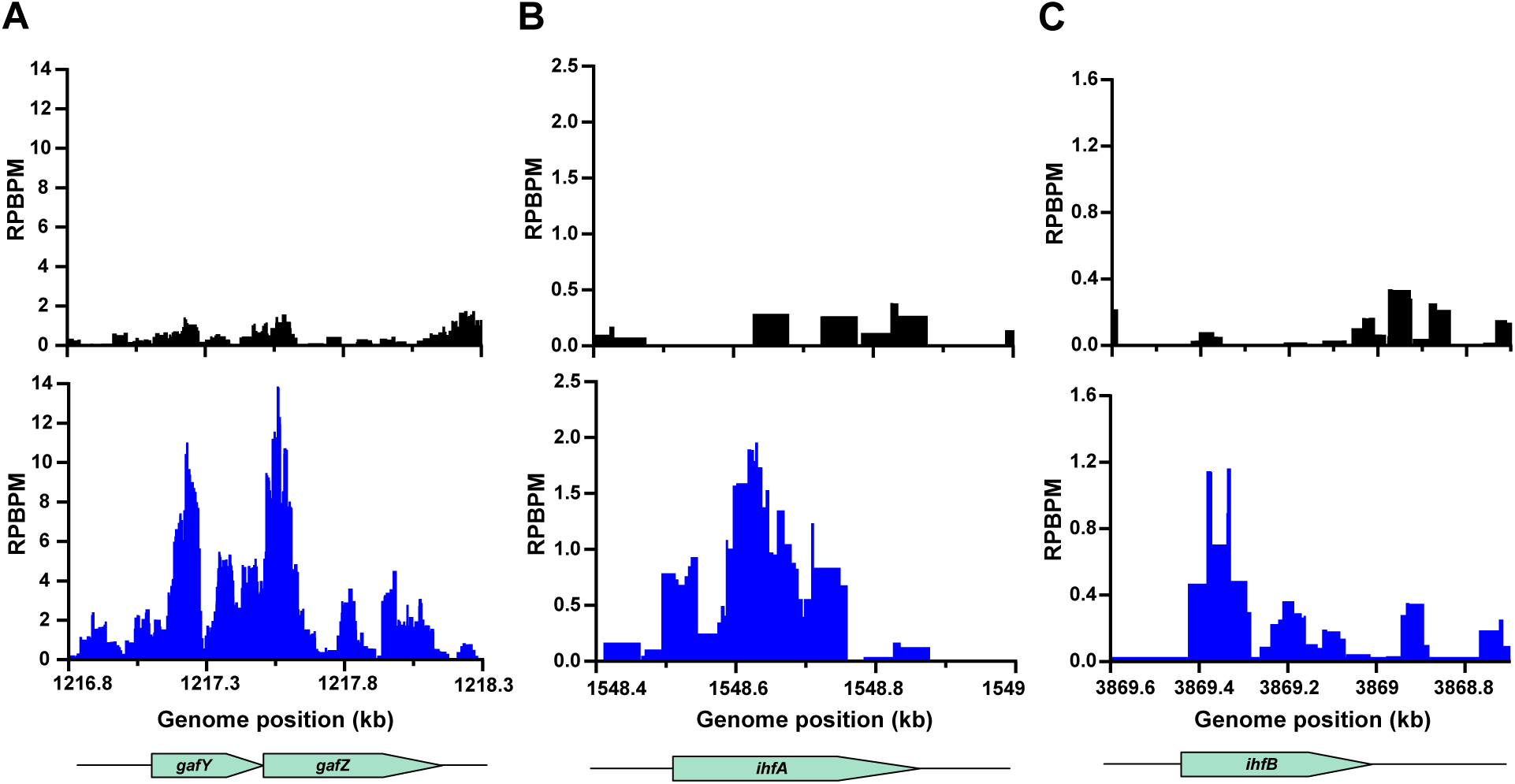
Transposon insertion frequency is greater within known GTA-activating genes in the Δ*rogA* mutant. Plots from the Tn-seq experiment (Figure 2B) showing the reads per base pair per million reads (RPBPM) at the genomic loci of *gafYZ* **(A)**, *ihfA* **(B)**, and *ihfB* **(C)** indicating the higher frequency of transposon insertions in these genes within the Δ*rogA* strain (bottom, blue trace) compared to the wildtype strain (top, black trace). Three independent repeats were performed.

**Figure S3.**
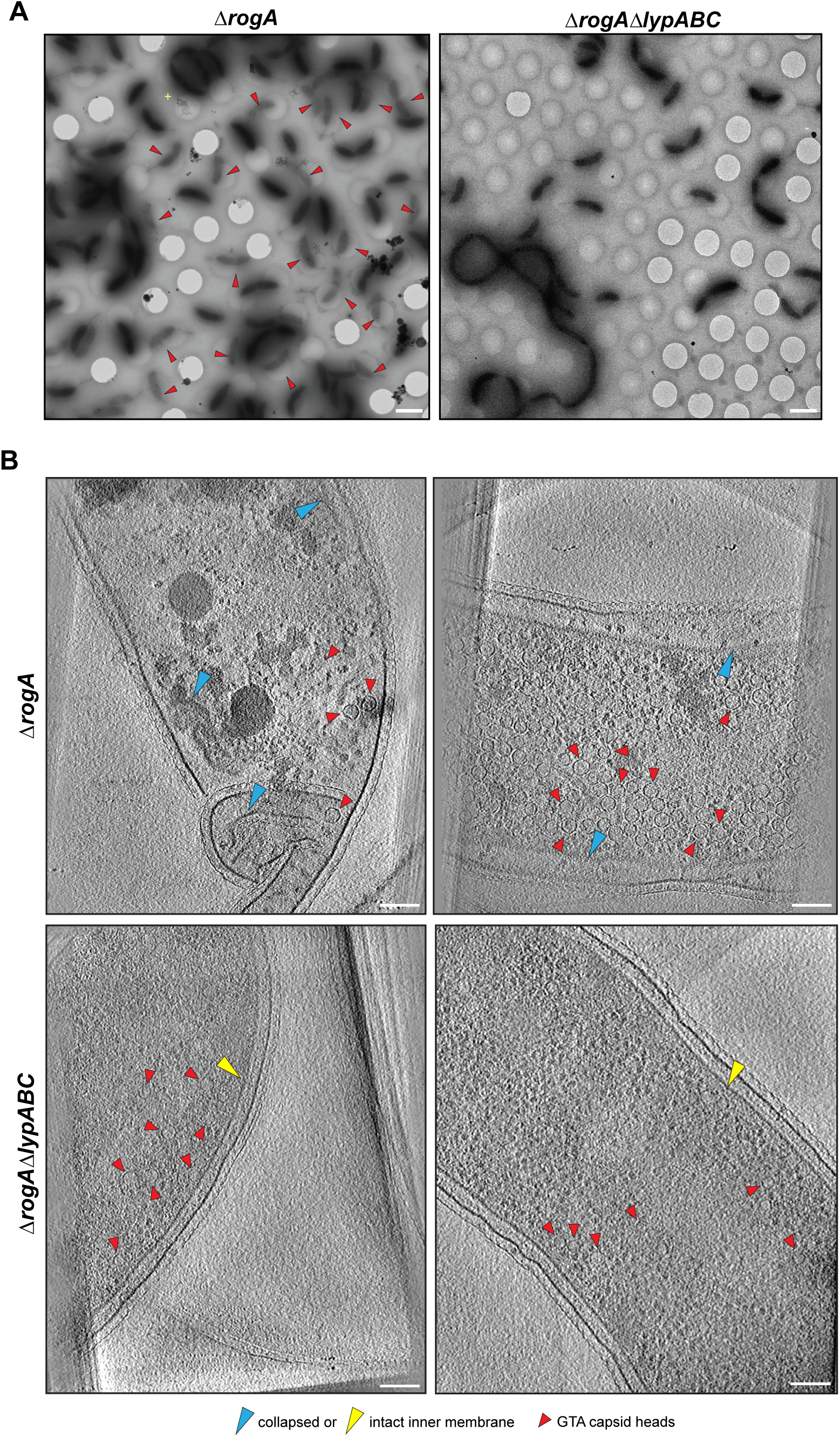
Cryo-electron microscopy and tomography of GTA-producing strains. **(A)** Cryo-electron microscopy search images from either the Δ*rogA* strain (left) or Δ*rogAΔlypABC* strain (right). Red arrows indicate lysing ghost cells in the Δ*rogA* strain only. Scale bar: 2 µm. Images are representative of two independent repeats. **(B)** Additional cryo-electron tomograms of Δ*rogA* (top row) and Δ*rogA*Δ*lypABC* (bottom row) cells to supplement Figure 3C. Red arrows: GTA capsid heads, blue arrow: collapsed inner membrane, yellow arrow: intact inner membrane. Scale bar: 100 nm. Images are representative of two independent experiments.

**Figure S4.**
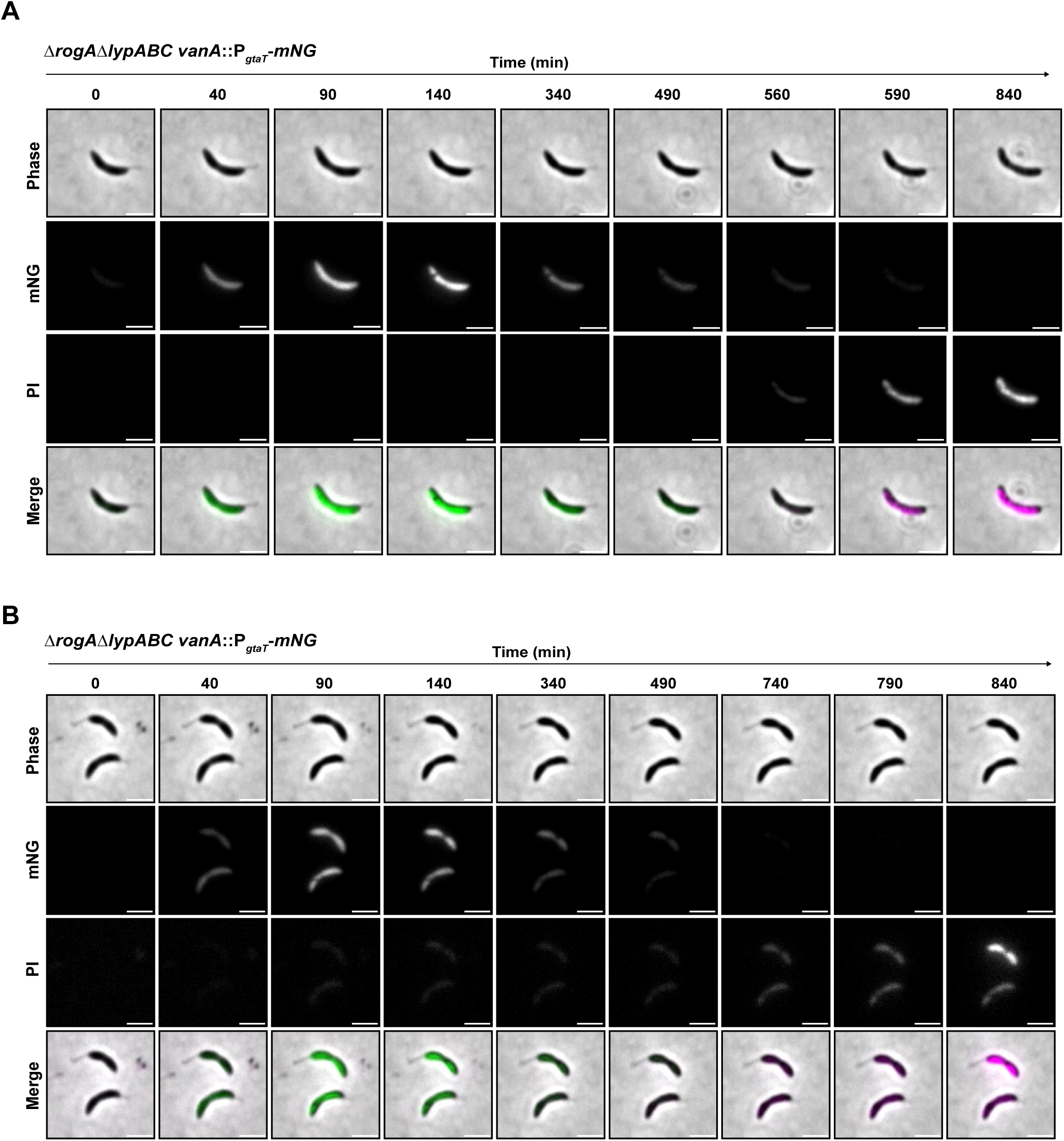
GTA-producing cells die but do not lyse in the absence of *lypABC*. Two examples of fluorescence microscopy images from time-lapse experiments performed with the Δ*rogA*Δ*lypABC vanA:*:P*_gtaT_* -*mNG* strain showing GTA activation (mNG signal), followed by cell death (PI signal) but no lysis or transition to a ghost cell. Time-lapses can be viewed in Movie S4 **(A)** and Movie S5 **(B)**. Data are representative of at least three independent repeats. Scale bar: 2 µm.

**Figure S5.**
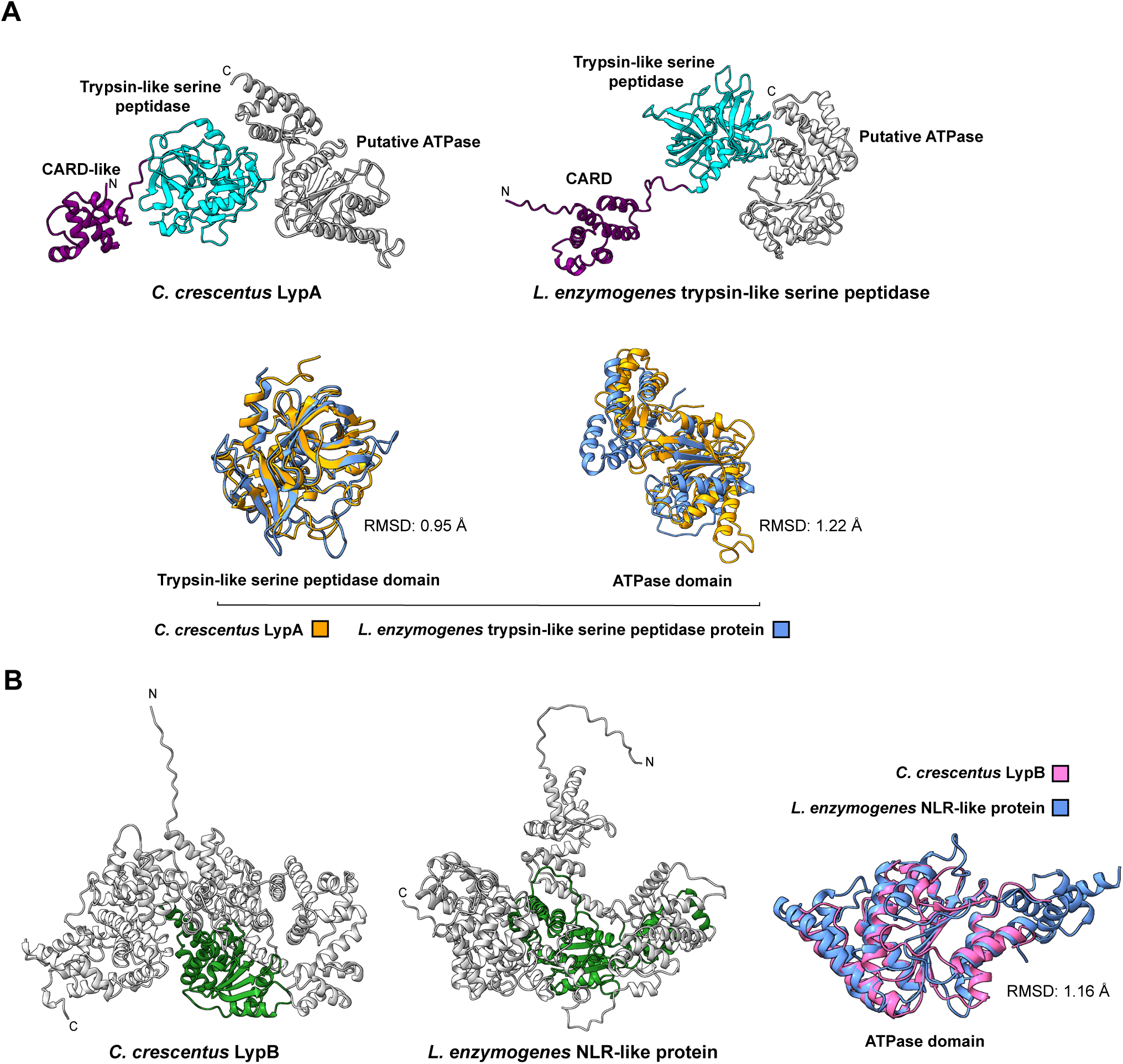
AlphaFold3 models of *C. crescentus* LypA and LypB compared to their *L. enzymogenes* CARD-NLR homologues. **(A)** Top: AlphaFold3 models of *C. crescentus* LypA (top left) and the *L. enzymogenes* CARD-NLR system trypsin-like serine peptidase protein (top right) coloured according to domain organisation. Bottom: AlphaFold3 model alignments of the trypsin-like serine peptidase domains (bottom left) and ATPase domains (bottom right) in isolation. Orange: *C. crescentus* LypA; blue: *L. enzymogenes* trypsin-like serine peptidase. Trypsin-like peptidase RMSD: 0.95 Å, ATPase RMSD: 1.22 Å. **(B)** AlphaFold3 models of *C. crescentus* LypB (left) and the *L. enzymogenes* CARD-NLR system NLR-like protein (middle) coloured according to domain organisation. Right: AlphaFold3 model alignment of the ATPase domains in isolation. Magenta: *C. crescentus* LypB; blue: *L. enzymogenes* NLR-like protein. RMSD: 1.16 Å.

**Figure S6.**
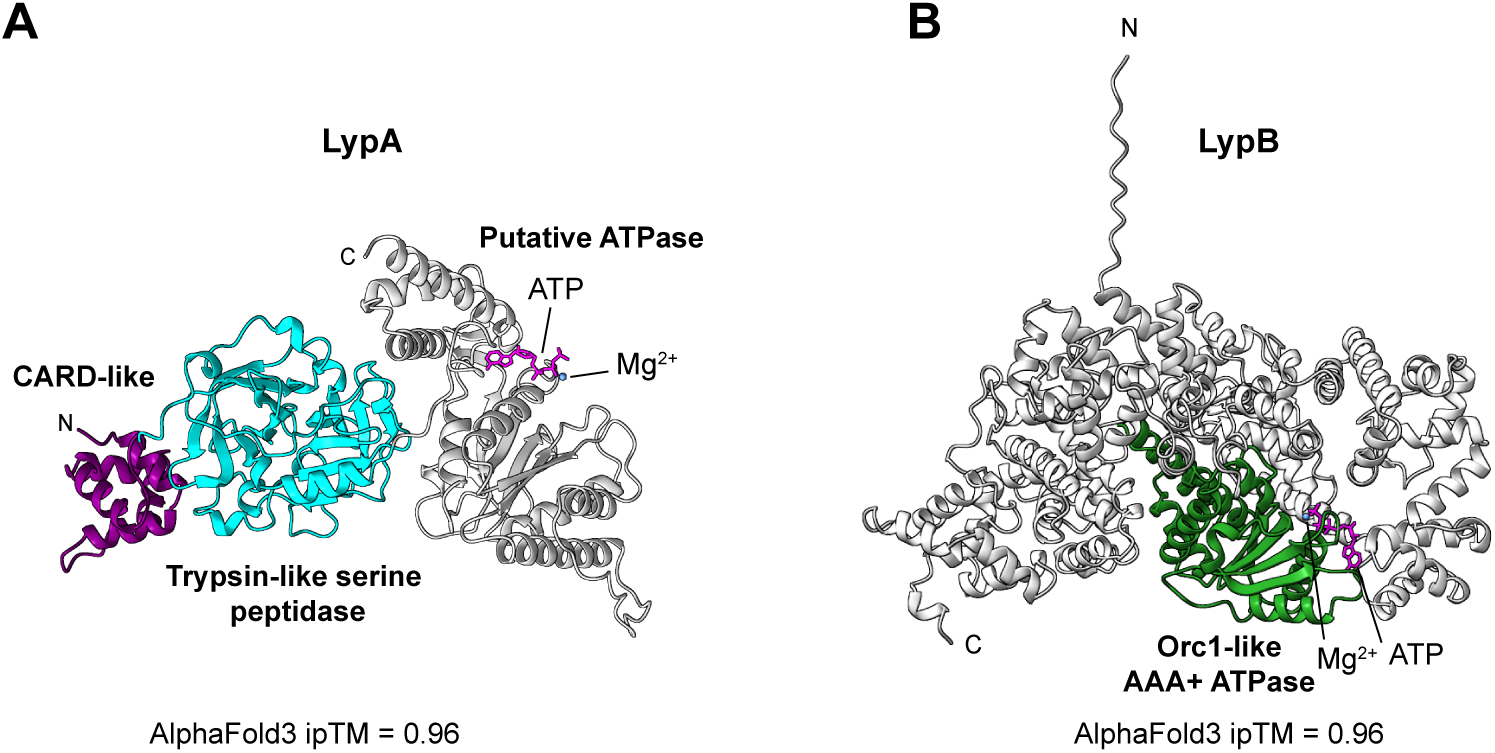
Structural predictions suggest LypA and LypB could potentially bind ATP. AlphaFold3 structural predictions of either LypA **(A)** or LypB **(B)** in complex with ATP (stick-format, magenta) and a magnesium Mg^2+^ ion (blue). The high ipTM score (0.96 for both) suggests that both LypA and LypB may be capable of binding ATP. Domains are coloured according to structures presented in Figure 4. The C-terminal domain of LypA contains an ATPase-like fold yet lacks conserved Walker motifs and is therefore labelled as a putative ATPase.

**Figure S7.**
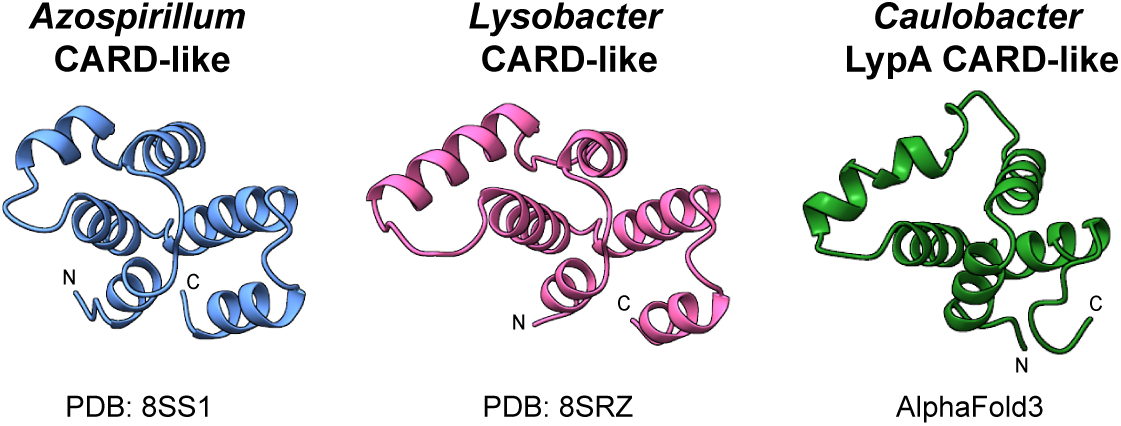
LypA contains a potential N-terminal CARD-like domain. Comparison of crystal structures of the CARD-like domains of proteins encoded by the CARD-NLR anti-phage defence systems of *Azospirillum* sp. TSO35-2 (left) and *Lysobacter enzymogenes* (centre) with the AlphaFold3-predicted structure of the N-terminal CARD-like domain (residues 1-80) of LypA from *C. crescentus* (right). PDB identifiers are shown beneath the crystal structures.

**Figure S8.**
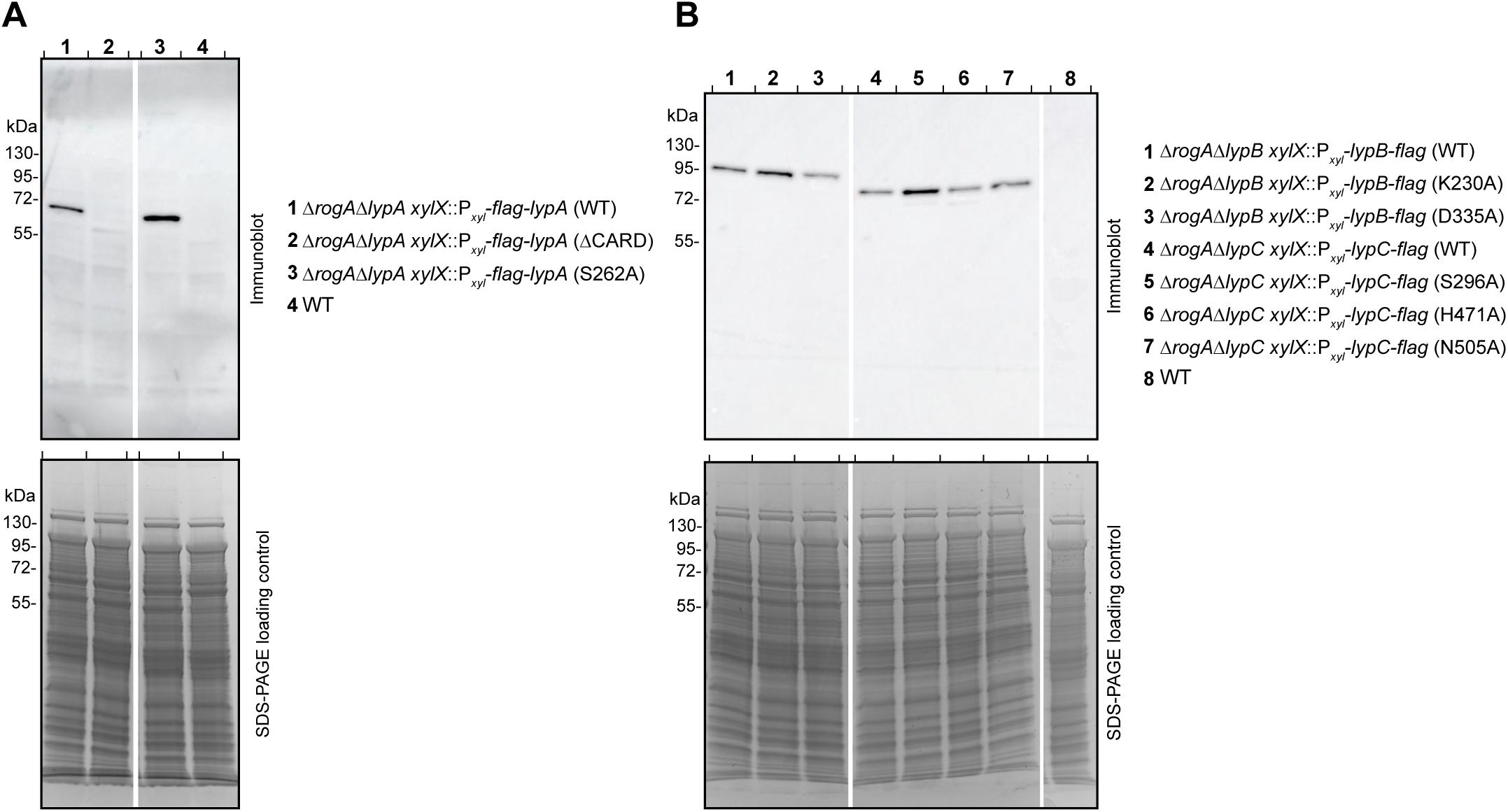
LypABC protein variants are stably produced *in vivo*. Immunoblots of total cell lysates from *C. crescentus* using a monoclonal anti-FLAG antibody showing stable production of either LypA protein variants **(A)** (with the exception of the ΔCARD mutant) or LypB and LypC protein variants **(B)** tested in Figure 4. The expected sizes for LypA, LypB and LypC are 65, 112, and 73 kDa, respectively. Separate Coomassie-stained SDS-PAGE gels were loaded with equal sample volumes to serve as loading controls. Immunoblots are representative of at least two independent experiments.

**Figure S9.**
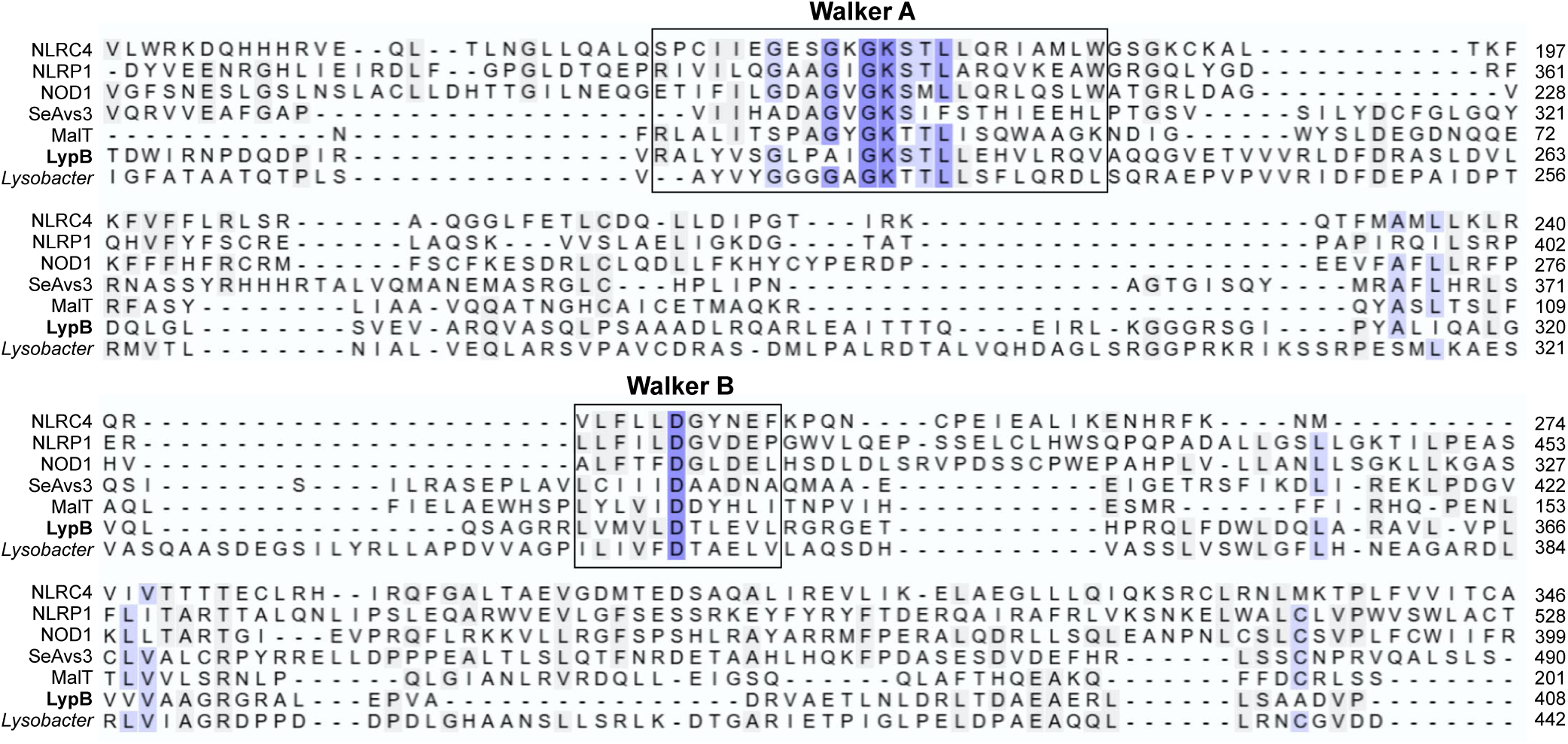
Multiple sequence alignment of the nucleotide binding domain of NLR and NLR-like ATPases including LypB. Sequence alignment of nucleotide binding domains from NLRC4 (*H. sapiens*, NCBI accession: AAH31555), NLRP1 (*H. sapiens*, NCBI accession: AAG30288), NOD1 (*H. sapiens*, NCBI accession: AAD29125), SeAvs3 (*Salmonella*, NCBI accession: WP_126523998.1), MalT (*E. coli*, NCBI accession: AAA8388.1), LypB (*C. crescentus*, NCBI accession: ACL94045.1), and *Lysobacter* (*L. enzymogenes*, NCBI accession: ALN60087.1). The Walker A and Walker B motifs are shown encased in black boxes.

**Figure S10.**
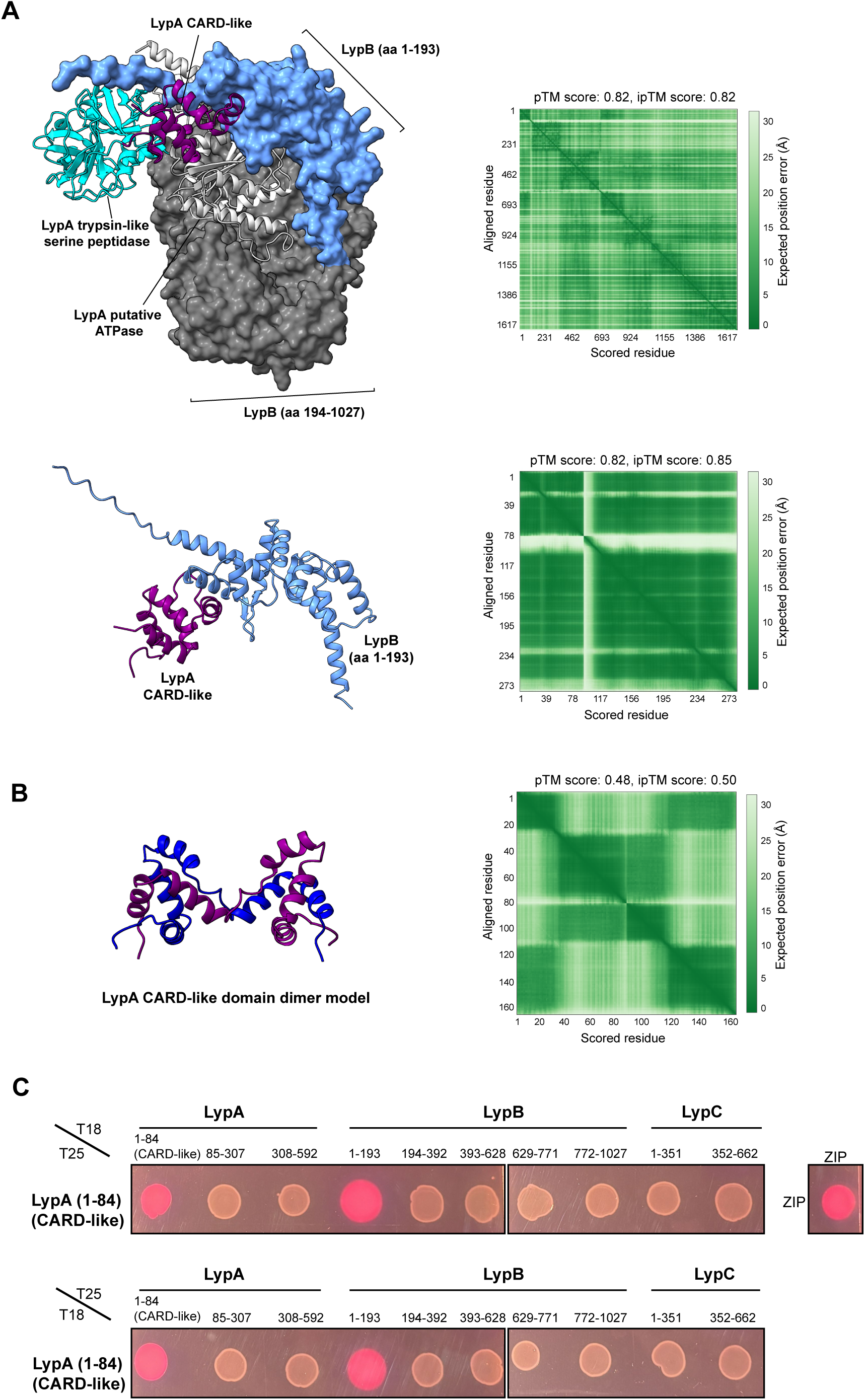
LypA and LypB interact via the CARD-like domain of LypA. **(A)** AlphaFold3-Multimer models of LypA and LypB (top) and the CARD-like domain of LypA with the N-terminal domain of LypB (aa 1-193) (bottom). Predicted aligned error plots are shown to the right of each model with model confidence scores stated above each error plot. The CARD-like domain, trypsin-like serine peptidase domain and putative ATPase domains of LypA are coloured purple, cyan, and light grey, respectively and depicted in ribbon format. LypB is either depicted in surface solid format (top) or in ribbon format (bottom). LypB aa 1-193: blue; aa 194-1027: dark grey. **(B)** AlphaFold3-Multimer model of the LypA CARD-like domain as a dimer (left) with the predicted aligned error plot and model confidence scores (right). **(C)** Bacterial two-hybrid assay showing interactions (red) or non-interactions (white) of the LypA CARD-like domain (aa 1-84) with each domain of LypA, LypB, and LypC individually. The amino acid range of each tested domain is shown above the spot plate images. ZIP-ZIP: positive interaction control. Proteins were tested in both T18 and T25 vectors (top and bottom rows). Three independent experiments were performed and representative data shown.

**Figure S11.**
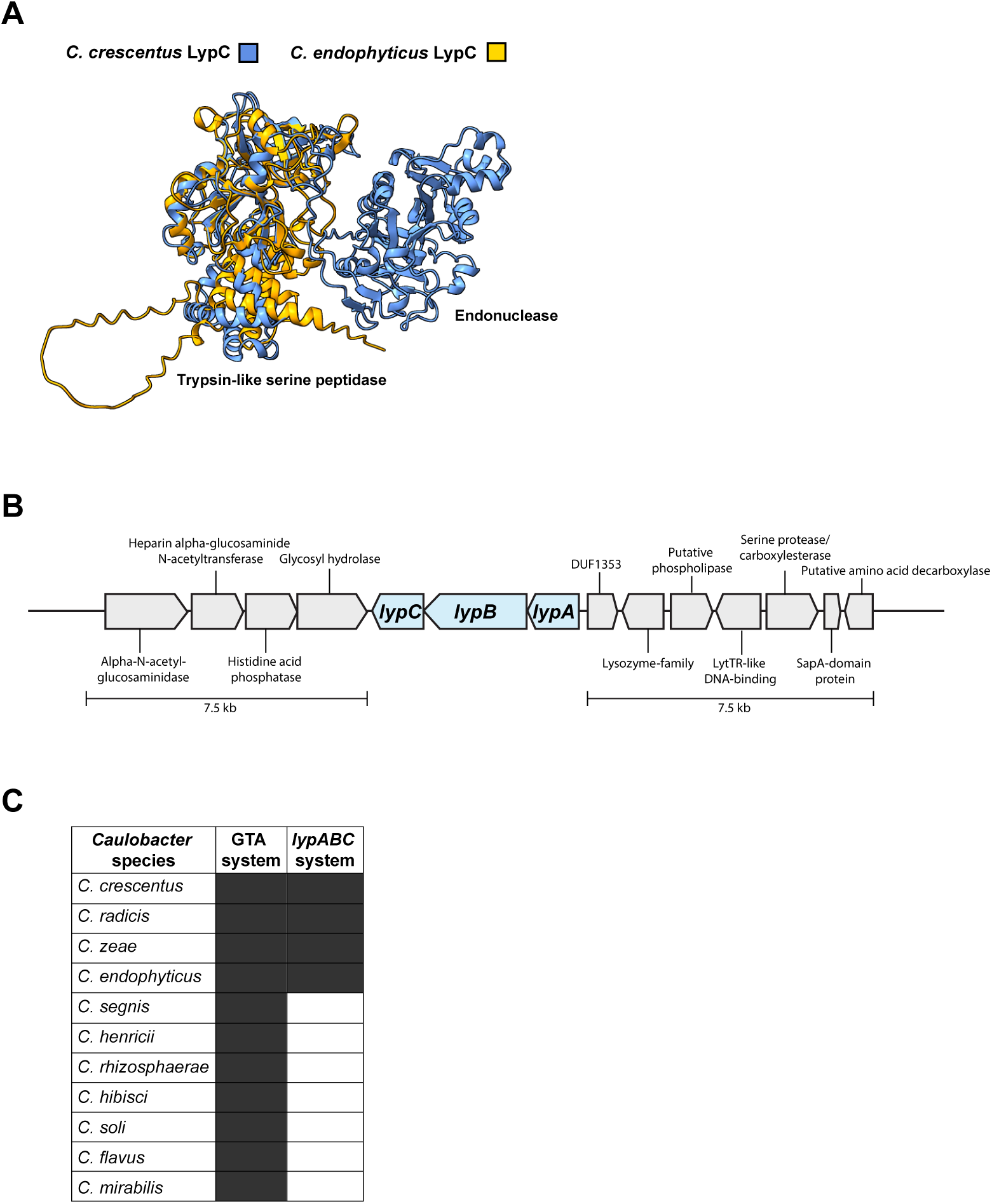
Genomic context and conservation of LypABC. **(A)** AlphaFold3 structural predictions for LypC from either *C. crescentus* (blue) or *C. endophyticus* (orange), highlighting the absence of a predicted C-terminal endonuclease domain in *C. endophyticus*. **(B)** Schematic of the genomic context of *lypABC* within *C. crescentus* strain NA1000. The genes are not encoded within a defence island like many anti-phage defence systems. The putative functions of genes flanking *lypABC* are annotated. **(C)** Table showing the predicted conservation of a GTA system and a complete *lypABC* system within different *Caulobacter* species (dark grey: present; white: absent). Only four species encode both systems.

**Figure S12.**
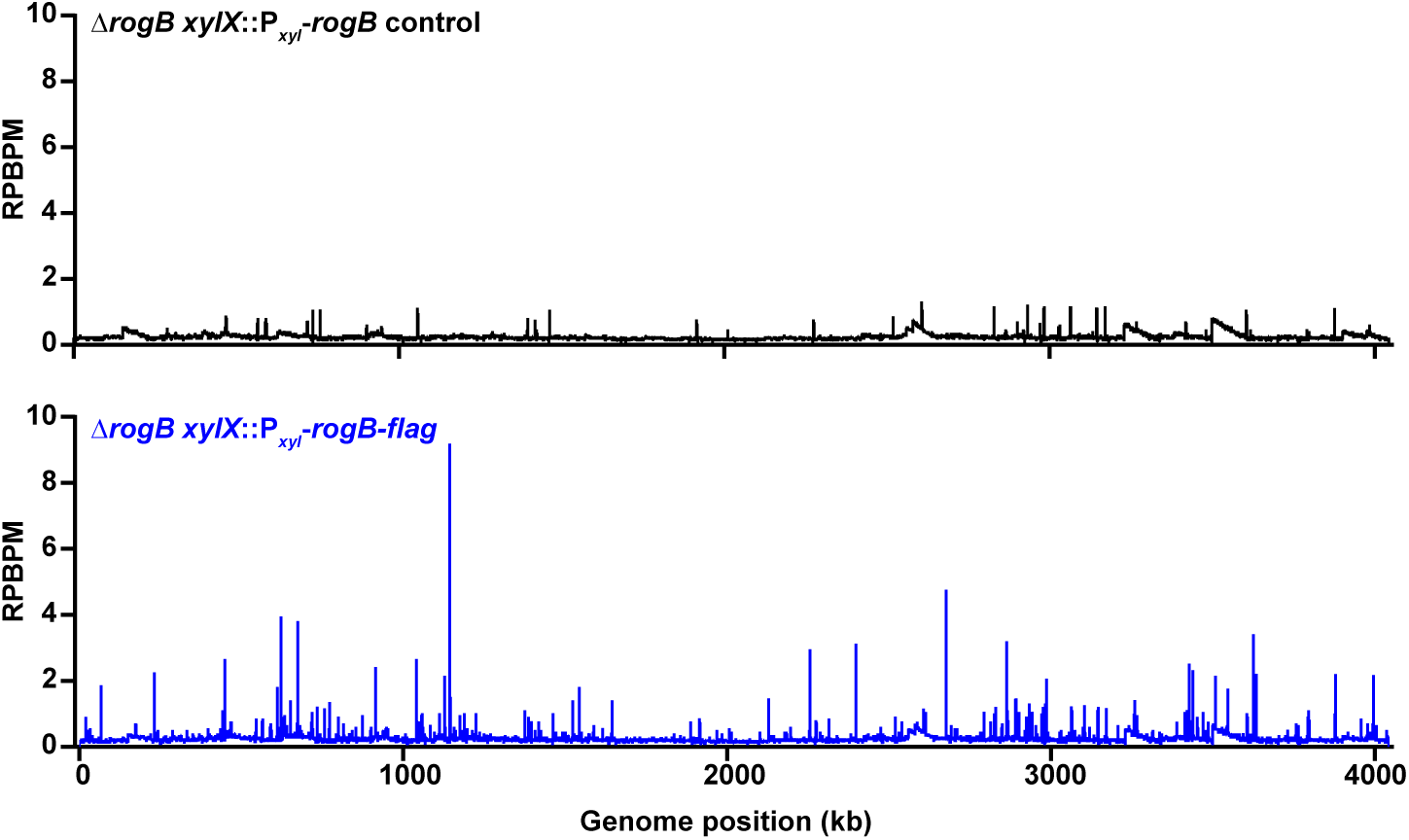
RogB binds to many sites on the *Caulobacter crescentus* genome. ChIP-seq profiles showing binding of RogB protein across the entire *C. crescentus* NA1000 genome *in vivo*. A *rogB-flag* (bottom profile, blue line) or untagged *rogB* (top profile, black line) allele was expressed from the xylose promoter (P*_xyl_*) in a Δ*rogB* mutant. The x-axis shows genomic position (kb) and the y-axis shows reads per base pair per million reads (RPBPM). Two independent repeats were performed and representative profiles are shown.

**Figure S13.**
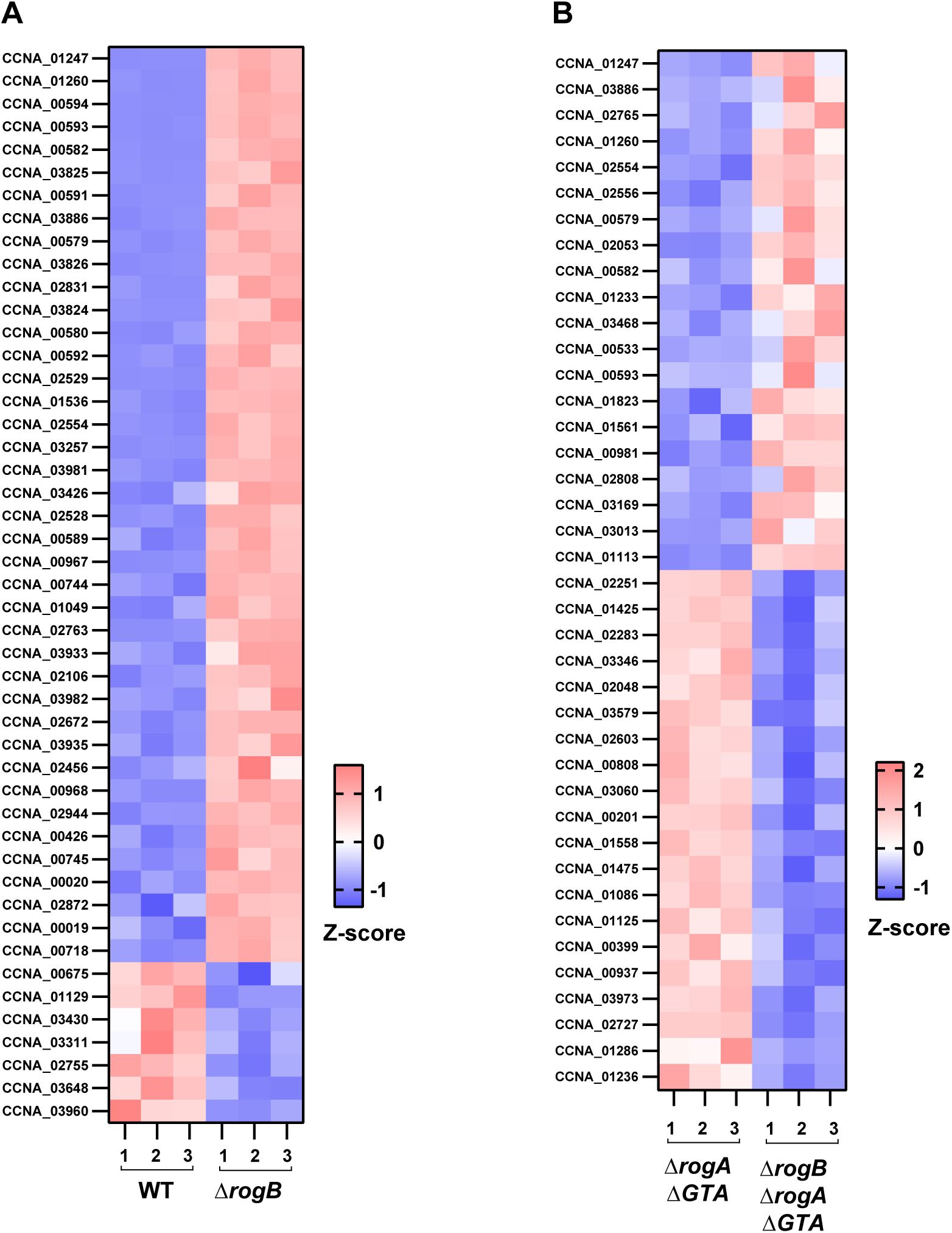
RNA-seq experiment heatmaps of top differentially expressed genes. Heatmaps showing the top differentially expressed genes for the RNA-seq experiments presented in Figure 6E (**A)** and Figure 7C **(B)**. Normalised read counts for each of three replicates (1-3) were used to generate Z-scores to show the differential expression for each gene across replicates in each mutant background. Red or blue values indicate higher or lower expression, respectively.

