## Supplementary tables, movies, materials and methods for "A bacterial CARD-NLR immune system controls the release of gene transfer agents": EJB_LypABC_manuscript_SI.docx

**Description of Supplementary Files**

**Supplementary materials and methods**

**Table S1** – DESeq2 results from Tn-seq experiments comparing Δ*rogA* with the wildtype strain

**Table S2** – MACS2 peaks called from ChIP-seq experiments of Δ*rogB xylX*:: P_xyl_-*rogB*-*flag* compared to the Δ*rogB xylX*::P_xyl_-*rogB* negative control

**Table S3** – DESeq2 results from RNA-seq experiments comparing Δ*rogB* with the wildtype strain

**Table S4** – MACS2 peaks called from ChIP-seq experiments of Δ*rogA*Δ*rogB xylX*::P_xyl_-*rogB*-*flag* compared to the Δ*rogA*Δ*rogB xylX*::P_xyl_-*rogB* negative control

**Table S5** – DESeq2 results from RNA-seq experiments comparing Δ*rogA*Δ*rogB*Δ*GTA* with the Δ*rogA*Δ*GTA* strain

**Table S6** – Strains constructed and used in this work

**Table S7** – Plasmids constructed and used in this work

**Table S8** – Oligonucleotides used in this work

**Movie S1** – Time-lapse movie of the *C. crescentus* Δ*rogA* *vanA*::P*_gtaT_-mN* fluorescent reporter strain

**Movie S2** – Time-lapse movie of the *C. crescentus* Δ*rogA* *vanA*::P*_gtaT_-mN* fluorescent reporter strain

**Movie S3** – Time-lapse movie of the *C. crescentus* Δ*rogA* *vanA*::P*_gtaT_-mN* fluorescent reporter strain

**Movie S4** – Time-lapse movie of the *C. crescentus* Δ*rogA* Δ*lypABC* *vanA*::P*_gtaT_-mN* fluorescent reporter strain growing in the presence of propidium iodide

**Movie S5** – Time-lapse movie of the *C. crescentus* Δ*rogA* Δ*lypABC* *vanA*::P*_gtaT_-mN* fluorescent reporter strain growing in the presence of propidium iodide

**Supplementary materials and methods**

**Construction of plasmids and strains**

**pET-21b::*rogB-6xhis***
The codon-optimised wildtype *rogB* gene with a 6x-*his* tag at the C-terminus was ordered as a double-stranded gBlock gene fragment (IDT) and cloned into NdeI + HindIII-cut pET-21b vector using 2x Gibson assembly master mix. Gibson reactions were introduced into chemically competent *E. coli* DH5α cells. A plasmid containing the correct insert was verified by Sanger sequencing and used to transform chemically competent *E. coli Rosetta* (DE3) cells, generating the strain Rosetta (DE3)::*rogB*-*his*.

**pXCHYC-2::*rogA***
The *rogA* gene was amplified from wildtype *C. crescentus* genomic DNA using EJB_005 + EJB_006 primers. PCR products were gel purified and cloned into NdeI + NheI-cut pXCHYC-2 integrative vector using 2x Gibson assembly master mix. Gibson reactions were introduced into chemically competent *E. coli* DH5α cells. A plasmid containing the correct insert was verified by Sanger sequencing and used to transform electrocompetent *C. crescentus* NA1000 Δ*rogA* cells to generate the strain Δ*rogA* *xylX*::P*_xyl_*-*rogA*.

**pVCHYC-1::P*_gtaT_-mNG***
The promoter region of the core GTA gene cluster (P*_gtaT_*) gene was amplified from wildtype *C. crescentus* genomic DNA using EJB_001 + EJB_002 primers. The *mNG* gene was amplified from plasmid template using EJB_003 + EJB_004 primers. PCR products were gel purified and cloned into NdeI + NheI-cut pVCHYC-1 integrative vector using 2x Gibson assembly master mix. Gibson reactions were introduced into chemically competent *E. coli* DH5α cells. A plasmid containing the correct insert was verified by Sanger sequencing and used to transform electrocompetent *C. crescentus* NA1000 cells to generate the strain *vanA*::P*_gtaT_-mNG*. Phage ΦCr30 transduction to delete *rogA* within this strain was performed as described previously^1^, generating the strain Δ*rogA* *vanA*::P*_gtaT_-mNG*.

**pNPTS138::Δ*lypABC***
Upstream and downstream regions (each 500 bp) flanking the *lypABC* genomic locus were amplified by PCR using EJB_007 + EJB_008 primers and EJB_009 + EJB_010 primers, respectively. PCR products were gel purified and cloned into BamHI + HindIII-cut pNPTS138 suicide vector using 2x Gibson assembly master mix. Gibson reactions were introduced into chemically competent *E. coli* DH5α cells. A plasmid containing the correct insert was verified by Sanger sequencing and used to transform electrocompetent *C. crescentus* NA1000 wildtype cells and delete *lypABC*. Phage transduction was used to delete *rogA* within this strain, generating the strain Δ*rogA*Δ*lypABC*. To construct the strain Δ*rogA*Δ*lypABC* *vanA*::P*_gtaT_*-*mNG*, the plasmid pVCHYC-1::P*_gtaT_-mNG* was first introduced into Δ*lypABC* followed by phage transduction to delete *rogA*.

**pNPTS138::Δ*lypA***
Upstream and downstream regions (each 500 bp) flanking the *lypA* gene were amplified by PCR using EJB_011 + EJB_012 primers and EJB_013 + EJB_014 primers, respectively. PCR products were gel purified and cloned into BamHI + HindIII-cut pNPTS138 suicide vector using 2x Gibson assembly master mix. Gibson reactions were introduced into chemically competent *E. coli* DH5α cells. A plasmid containing the correct insert was verified by Sanger sequencing and used to transform electrocompetent *C. crescentus* NA1000 wildtype cells and delete *lypA*. Phage transduction was used to delete *rogA* within this strain, generating the strain Δ*rogA*Δ*lypA*.

**pNPTS138::Δ*lypB***
Upstream and downstream regions (each 500 bp) flanking a region internal to the *lypB* gene (aa 230-335) were amplified by PCR using EJB_015 + EJB_016 primers and EJB_017 + EJB_018 primers, respectively. PCR products were gel purified and cloned into BamHI + HindIII-cut pNPTS138 suicide vector using 2x Gibson assembly master mix. Gibson reactions were introduced into chemically competent *E. coli* DH5α cells. A plasmid containing the correct insert was verified by Sanger sequencing and used to transform electrocompetent *C. crescentus* NA1000 wildtype cells and delete *lypB*. Phage transduction was used to delete *rogA* within this strain, generating the strain Δ*rogA*Δ*lypB*.

**pNPTS138::Δ*lypC***
Upstream and downstream regions (each 500 bp) flanking the *lypC* gene were amplified by PCR using EJB_019 + EJB_020 primers and EJB_021 + EJB_022 primers, respectively. PCR products were gel purified and cloned into BamHI + HindIII-cut pNPTS138 suicide vector using 2x Gibson assembly master mix. Gibson reactions were introduced into chemically competent *E. coli* DH5α cells. A plasmid containing the correct insert was verified by Sanger sequencing and used to transform electrocompetent *C. crescentus* NA1000 wildtype cells and delete *lypC*. Phage transduction was used to delete *rogA* within this strain, generating the strain Δ*rogA*Δ*lypC*.

**pXCHYC-2::*lypA***
The *lypA* gene was amplified from wildtype *C. crescentus* genomic DNA using EJB_023 + EJB_024 primers. PCR products were gel purified and cloned into NdeI + NheI-cut pXCHYC-2 integrative vector using 2x Gibson assembly master mix. Gibson reactions were introduced into chemically competent *E. coli* DH5α cells. A plasmid containing the correct insert was verified by Sanger sequencing and used to transform electrocompetent *C. crescentus* NA1000 Δ*rogA*Δ*lypA* cells to generate the strain Δ*rogA*Δ*lypA xylX*::P*_xyl_*-*lypA*.

**pXCHYC-2::*lypB***
The *lypB* gene was amplified from wildtype *C. crescentus* genomic DNA using EJB_025 + EJB_026 primers. PCR products were gel purified and cloned into NdeI + NheI-cut pXCHYC-2 integrative vector using 2x Gibson assembly master mix. Gibson reactions were introduced into chemically competent *E. coli* DH5α cells. A plasmid containing the correct insert was verified by Sanger sequencing and used to transform electrocompetent *C. crescentus* NA1000 Δ*rogA*Δ*lypB* cells to generate the strain Δ*rogA*Δ*lypB xylX*::P*_xyl_*-*lypB*.

**pXCHYC-2::*lypC***
The *lypC* gene was amplified from wildtype *C. crescentus* genomic DNA using EJB_027 + EJB_028 primers. PCR products were gel purified and cloned into NdeI + NheI-cut pXCHYC-2 integrative vector using 2x Gibson assembly master mix. Gibson reactions were introduced into chemically competent *E. coli* DH5α cells. A plasmid containing the correct insert was verified by Sanger sequencing and used to transform electrocompetent *C. crescentus* NA1000 Δ*rogA*Δ*lypC* cells to generate the strain Δ*rogA*Δ*lypC xylX*::P*_xyl_*-*lypC*.

**pXCHYC-2::*flag-lypA* (WT)**
The codon-optimised wildtype *lypA* gene with a 1x-*flag* tag at the N-terminus was ordered as a gBlock and cloned into NdeI + NheI-cut pXCHYC-2 integrative vector using 2x Gibson assembly master mix. Gibson reactions were introduced into chemically competent *E. coli* DH5α cells. A plasmid containing the correct insert was verified by Sanger sequencing and used to transform electrocompetent *C. crescentus* NA1000 Δ*rogA*Δ*lypA* cells, generating the strain Δ*rogA*Δ*lypA xylX*::P*_xyl_*-*flag*-*lypA*.

**pXCHYC-2::*flag-lypA* (ΔCARD)**
The codon-optimised *lypA* gene (containing a truncation of the aa 1-80 CARD-like domain) with a 1x-*flag* tag at the N-terminus was ordered as a gBlock and cloned into NdeI + NheI-cut pXCHYC-2 integrative vector using 2x Gibson assembly master mix. Gibson reactions were introduced into chemically competent *E. coli* DH5α cells. A plasmid containing the correct insert was verified by Sanger sequencing and used to transform electrocompetent *C. crescentus* NA1000 Δ*rogA*Δ*lypA* cells, generating the strain Δ*rogA*Δ*lypA xylX*::P*_xyl_*-*flag*-*lypA* (ΔCARD).

**pXCHYC-2::*flag-lypA* (S262A)**
The codon-optimised *lypA* gene (containing a mutation of S262A) with a 1x-*flag* tag at the N-terminus was ordered as a gBlock and cloned into NdeI + NheI-cut pXCHYC-2 integrative vector using 2x Gibson assembly master mix. Gibson reactions were introduced into chemically competent *E. coli* DH5α cells. A plasmid containing the correct insert was verified by Sanger sequencing and used to transform electrocompetent *C. crescentus* NA1000 Δ*rogA*Δ*lypA* cells, generating the strain Δ*rogA*Δ*lypA xylX*::P*_xyl_*-*flag*-*lypA* (S262A).

**pXCHYC-2::*lypB-flag* (WT)**
The codon-optimised wildtype *lypB* gene with a 1x-*flag* tag at the C-terminus was ordered as a gBlock and cloned into NdeI + NheI-cut pXCHYC-2 integrative vector using 2x Gibson assembly master mix. Gibson reactions were introduced into chemically competent *E. coli* DH5α cells. A plasmid containing the correct insert was verified by Sanger sequencing and used to transform electrocompetent *C. crescentus* NA1000 Δ*rogA*Δ*lypB* cells, generating the strain Δ*rogA*Δ*lypB xylX*::P*_xyl_*-*lypB-flag*.

**pXCHYC-2::*lypB-flag* (K230A)**
The codon-optimised *lypB* gene (containing a mutation of K230A) with a 1x-*flag* tag at the C-terminus was ordered as a gBlock and cloned into NdeI + NheI-cut pXCHYC-2 integrative vector using 2x Gibson assembly master mix. Gibson reactions were introduced into chemically competent *E. coli* DH5α cells. A plasmid containing the correct insert was verified by Sanger sequencing and used to transform electrocompetent *C. crescentus* NA1000 Δ*rogA*Δ*lypB* cells, generating the strain Δ*rogA*Δ*lypB xylX*::P*_xyl_*-*lypB-flag* (K230A).

**pXCHYC-2::*lypB-flag* (D335A)**
The codon-optimised *lypB* gene (containing a mutation of D335A) with a 1x-*flag* tag at the C-terminus was ordered as a gBlock and cloned into NdeI + NheI-cut pXCHYC-2 integrative vector using 2x Gibson assembly master mix. Gibson reactions were introduced into chemically competent *E. coli* DH5α cells. A plasmid containing the correct insert was verified by Sanger sequencing and used to transform electrocompetent *C. crescentus* NA1000 Δ*rogA*Δ*lypB* cells, generating the strain Δ*rogA*Δ*lypB xylX*::P*_xyl_*-*lypB-flag* (D335A).

**pXCHYC-2::*lypC-flag* (WT)**
The codon-optimised wildtype *lypC* gene with a 1x-*flag* tag at the C-terminus was ordered as a gBlock and cloned into NdeI + NheI-cut pXCHYC-2 integrative vector using 2x Gibson assembly master mix. Gibson reactions were introduced into chemically competent *E. coli* DH5α cells. A plasmid containing the correct insert was verified by Sanger sequencing and used to transform electrocompetent *C. crescentus* NA1000 Δ*rogA*Δ*lypC* cells, generating the strain Δ*rogA*Δ*lypB xylX*::P*_xyl_*-*lypC-flag*.

**pXCHYC-2::*lypC-flag* (S296A)**
The codon-optimised *lypC* gene (containing a mutation of S296A) with a 1x-*flag* tag at the C-terminus was ordered as a gBlock and cloned into NdeI + NheI-cut pXCHYC-2 integrative vector using 2x Gibson assembly master mix. Gibson reactions were introduced into chemically competent *E. coli* DH5α cells. A plasmid containing the correct insert was verified by Sanger sequencing and used to transform electrocompetent *C. crescentus* NA1000 Δ*rogA*Δ*lypC* cells, generating the strain Δ*rogA*Δ*lypB xylX*::P*_xyl_*-*lypC-flag* (S296A).

**pXCHYC-2::*lypC-flag* (H471A)**
The codon-optimised *lypC* gene (containing a mutation of H471A) with a 1x-*flag* tag at the C-terminus was ordered as a gBlock and cloned into NdeI + NheI-cut pXCHYC-2 integrative vector using 2x Gibson assembly master mix. Gibson reactions were introduced into chemically competent *E. coli* DH5α cells. A plasmid containing the correct insert was verified by Sanger sequencing and used to transform electrocompetent *C. crescentus* NA1000 Δ*rogA*Δ*lypC* cells, generating the strain Δ*rogA*Δ*lypB xylX*::P*_xyl_*-*lypC-flag* (H471A).

**pXCHYC-2::*lypC-flag* (N505A)**
The codon-optimised *lypC* gene (containing a mutation of N505A) with a 1x-*flag* tag at the C-terminus was ordered as a gBlock and cloned into NdeI + NheI-cut pXCHYC-2 integrative vector using 2x Gibson assembly master mix. Gibson reactions were introduced into chemically competent *E. coli* DH5α cells. A plasmid containing the correct insert was verified by Sanger sequencing and used to transform electrocompetent *C. crescentus* NA1000 Δ*rogA*Δ*lypC* cells, generating the strain Δ*rogA*Δ*lypB xylX*::P*_xyl_*-*lypC-flag* (N505A).

**pNPTS::P*_cumate_-lypABC***
Upstream and downstream regions (each 500 bp) flanking the start codon of the *lypA* gene were amplified by PCR using EJB_029 + EJB_030 primers and EJB_033 + EJB_034 primers, respectively. The cumate promoter sequence was amplified from a plasmid template using primers EJB_031 and EJB_032. PCR products were gel purified and cloned into BamHI + HindIII-cut pNPTS138 suicide vector using 2x Gibson assembly master mix. Gibson reactions were introduced into chemically competent *E. coli* DH5α cells. A plasmid containing the correct insert was verified by Sanger sequencing and used to transform electrocompetent *C. crescentus* NA1000 wildtype cells*,* generating the strain P*_cumate_-lypABC.* Phage transduction was used to delete *rogA* within this strain, generating the strain Δ*rogA* P*_cumate_-lypABC*.

**pNPTS::Δ*rogB***
Upstream and downstream regions (each 500 bp) flanking the *rogB* gene were amplified by PCR using EJB_035 + EJB_036 primers and EJB_037 + EJB_038 primers, respectively. PCR products were gel purified and cloned into BamHI + HindIII-cut pNPTS138 suicide vector using 2x Gibson assembly master mix. Gibson reactions were introduced into chemically competent *E. coli* DH5α cells. A plasmid containing the correct insert was verified by Sanger sequencing and used to transform electrocompetent *C. crescentus* NA1000 wildtype cells and delete *rogB,* generating the strain Δ*rogB.* Phage transduction was used to delete *rogA* within this strain, generating the strain Δ*rogA*Δ*rogB*.

**pXCHYC-2::*rogB-flag***
The *rogB* gene was amplified from wildtype *C. crescentus* genomic DNA using EJB_039 + EJB_040 primers which contained a 1x-*flag* tag. PCR products were gel purified and cloned into NdeI + NheI-cut pXCHYC-2 integrative vector using 2x Gibson assembly master mix. Gibson reactions were introduced into chemically competent *E. coli* DH5α cells. A plasmid containing the correct insert was verified by Sanger sequencing and used to transform electrocompetent *C. crescentus* NA1000 Δ*rogB* cells to generate the strain Δ*rogB xylX*::P*_xyl_*-*rogB-flag*. Phage transduction was used to delete *rogA* within this strain, generating the strain Δ*rogA*Δ*rogB xylX*::P*_xyl_*-*rogB-flag*.

**pXCHYC-2::*rogB***
The *rogB* gene was amplified from wildtype *C. crescentus* genomic DNA using EJB_041 + EJB_042 primers. PCR products were gel purified and cloned into NdeI + NheI-cut pXCHYC-2 integrative vector using 2x Gibson assembly master mix. Gibson reactions were introduced into chemically competent *E. coli* DH5α cells. A plasmid containing the correct insert was verified by Sanger sequencing and used to transform electrocompetent *C. crescentus* NA1000 Δ*rogB* cells to generate the strain Δ*rogB xylX*::P*_xyl_*-*rogB*. Phage transduction was used to delete *rogA* within this strain, generating the strain Δ*rogA*Δ*rogB xylX*::P*_xyl_*-*rogB*.

**pXCHYC-2::P*_rogB_-rogB***
The *rogB* gene including its native promoter (P*_rogB_*) was amplified from wildtype *C. crescentus* genomic DNA using EJB_043 + EJB_044 primers. PCR products were gel purified and cloned into NdeI + NheI-cut pXCHYC-2 integrative vector using 2x Gibson assembly master mix. Gibson reactions were introduced into chemically competent *E. coli* DH5α cells. A plasmid containing the correct insert was verified by Sanger sequencing and used to transform electrocompetent *C. crescentus* NA1000 Δ*rogB* cells to generate the strain Δ*rogB xylX*::P*_rogB_*-*rogB*. Phage transduction was used to delete *rogA* within this strain, generating the strain Δ*rogA*Δ*rogB xylX*::P*_rogB_*-*rogB*.

**pNPTS::Δ*GTA***
Upstream and downstream regions (each 500 bp) flanking the core GTA cluster (CCNA_02861-CCNA_02880) were amplified by PCR using EJB_045 + EJB_046 primers and EJB_047 + EJB_048 primers, respectively. PCR products were gel purified and cloned into BamHI + HindIII-cut pNPTS138 suicide vector using 2x Gibson assembly master mix. Gibson reactions were introduced into chemically competent *E. coli* DH5a cells. A plasmid containing the correct insert was verified by Sanger sequencing and used to transform electrocompetent *C. crescentus* NA1000 wildtype cells and delete the gene cluster*,* generating the strain Δ*GTA.* Phage transduction was used to delete *rogA* within this strain, generating the strain Δ*rogA*Δ*GTA*. To construct the strain Δ*rogB*Δ*rogA*Δ*GTA*, *rogB* was first deleted within the Δ*GTA* strain to make Δ*rogB*Δ*GTA* and then phage transduction was performed to delete *rogA*.

**pKT25::*lypA/lypB/lypC* (individual domain) and pUT18C::*lypA/lypB/lypC* (individual domain)**
DNA encoding individual domains of LypA, LypB, and LypC were chemically synthesised as double-stranded gBlock DNA fragments (IDT). These DNA fragments were assembled into a BamHI-EcoRI-cut pKT25 vector or a BamHI-EcoRI-cut pUT18C vector using 2x Gibson assembly master mix. The resulting plasmid was verified by whole-plasmid sequencing (Plasmidsaurus). To construct bacterial two-hybrid strains, chemically-competent BTH101 *E. coli cya^-^* cells were co-transformed with combinations of pKT25 and pUT18C plasmids above.
